# Prospective Fecal Microbiomics Biomarkers for Chronic Wasting Disease

**DOI:** 10.1101/2023.08.21.554213

**Authors:** Adam Didier, Maureen Bourner, Guy Kleks, Avihai Zolty, Brajendra Kumar, Tracy Nichols, Karie Durynski, Susan Bender, Michelle Gibison, Lisa Murphy, Julie C. Ellis, Dawei W. Dong, Anna Kashina

## Abstract

Chronic wasting disease (CWD) is a naturally occurring prion disease in cervids that has been rapidly proliferating in the US. Here we investigated a potential link between CWD infection and gut microbiome by analyzing 50 fecal samples obtained from CWD-positive animals of different sexes from various regions in the US, compared to 50 CWD-negative controls using high throughput sequencing of the 16S ribosomal RNA and targeted metabolomics. Our analysis reveals promising trends in the gut microbiota that could potentially be CWD-dependent, including several bacterial taxa at each rank level, as well as taxa pairs, that can differentiate between CWD-negative and CWD-positive deer. At each rank level, these taxa and taxa pairs could facilitate identification of around 70% of both the CWD-negative and the CWD-positive samples. Our results provide a potential tool for diagnostics and surveillance of CWD in the wild, as well as conceptual advances in our understanding of the disease.

**Importance:** This is a comprehensive study that tests the connection between the composition of the gut microbiome in deer in response to Chronic Wasting Disease (CWD). We analyzed 50 fecal samples obtained from CWD-positive animals compared to 50 CWD-negative controls to identify CWD-dependent changes in the gut microbiome, matched with the analysis of fecal metabolites. Our results show promising trends suggesting that fecal microbial composition can directly correspond to CWD disease status. These results point to microbial composition of the feces as a potential tool for diagnostics and surveillance of CWD in the wild, including non-invasive CWD detection in asymptomatic deer and deer habitats, and enable conceptual advances in our understanding of the disease.

## Introduction

Chronic wasting disease (CWD) is a naturally occurring infectious, fatal, transmissible spongiform encephalopathy of cervids. Environmental contamination and excreta (e.g., saliva, urine, and feces) are thought to play a pivotal role in the rapid proliferation of CWD across North America over the past five decades (*1*).

CWD belongs to a broader class of prion diseases, caused by accumulation of abnormally misfolded prion proteins in the animals’ tissues and organs (*2*). While the full spectrum of organismal effects of prion diseases is still being characterized, the most affected tissue is the brain, where misfolded prions cause neuronal loss that leads to progressive neuronal dysfunction and brain damage, which is eventually fatal. CWD is believed to be transmissible across species. Although CWD transmission to humans has not been directly demonstrated, such transmission has been found to occur in another prion disease (e.g., bovine spongiform encephalopathy or Mad Cow Disease), and thus it remains a concerning possibility. Importantly, ingestion of infectious prions represents an established route of infection, and therefore human consumption of CWD-infected meat is of strong concern. Thus, surveillance and diagnostics are very important to CWD prevention and control, and represents a global challenge for animal health.

Currently, disease confirmation in cervids relies largely on post-mortem detection of infectious prions in the medial retropharyngeal lymph nodes or obex in the brain via immunohistochemistry (IHC). Recently, a Real-Time Quaking-Induced Conversion (RT-QuIC) assay has been developed, which can detect CWD from deer ear punches with high sensitivity (*3*). Additional antemortem samples that can be tested by RT-QuIC include feces, urine, and rectoanal mucosa-associated lymphoid tissue (RAMALT) biopsies. However, despite the long-standing recognition of CWD and progress made in the understanding of the disease, there is no non-invasive live-animal test with sensitivity greater than or equal to that of postmortem IHC or ELISA (*4–6*). While recent studies propose the use of RT-QuIC for CWD detection in fecal samples (*7*), no methods have currently been sufficiently optimized to enable disease surveillance using animal-derived materials and by-products in the wild.

While routes of CWD propagation are still being investigated, healthy animals are believed to acquire this disease by oral exposure to infected animal by-products containing misfolded prions, including feces, saliva, urine, and animal remains, which are accidentally ingested by the deer in the contaminated environment, and are then absorbed in the oral cavity and the digestive tract (*8–14*). Thus, the gastrointestinal tract represents a major route of CWD infection and has a high potential of being affected by CWD.

Gastrointestinal health as well as normal animal physiology greatly depend on gut microbiota, a multi-species population of symbiotic and pathogenic microorganisms normally residing in the intestines of all animals. Gut microbiota respond to a wide variety of diseases and physiological changes in the body (see, e.g., (*15*)) and are essential for maintaining the normal gut health and barrier function (*16, 17*). Based on the available evidence, recent papers propose a direct link between the gut microbiota and the pathogenesis and pathology of prion diseases (*18–20*), but this highly promising field is still in its infancy.

Gut microbiota composition in the digestive tract (microbiome) can be analyzed using feces, which contain a representative sample of bacteria from each individual animal. Feces are commonly found in natural deer habitats and can be collected without disturbing the environment or the need to trap or handle animals. Thus, identification of potential feces-based diagnostic markers of CWD would provide a very useful tool for disease surveillance and control. Studies analyzing fecal samples from deer with CWD are emerging in the field (*7, 21–25*), but currently no robust biomarkers that can inform disease surveillance, diagnostics, and its effects on normal animal physiology have been established.

Here we used fecal samples obtained from 50 CWD-positive farmed white-tailed deer (*Odocoileus virginianus*) of different sexes from various locations in the US, as well as 50 CWD-negative farmed white-tailed deer controls, to identify CWD-dependent changes in the deer microbiome. Using high throughput sequencing of the 16S ribosomal RNA gene, we identified promising trends in the gut microbiota that could potentially be CWD-dependent. While, in agreement with previously published studies, geographical origin exerts strong influence on the microbial composition of the gut, using CWD as a variable reveals 25 bacterial taxa that are differentially abundant between control and CWD-positive deer in our samples and can be explored as potential markers of CWD. Furthermore, we performed targeted metabolomics on the same fecal samples and cross-omics correlative analysis of these data with the microbial composition of the samples, to identify potential changes in host and microbial metabolites. Collectively, these changes represent potential microbial signatures that may prove to be specific to CWD-infected animals. Longer term, these results point to a possibility of non-invasive diagnostics and surveillance of CWD in wild and farmed white-tailed deer using fecal samples, as well as conceptual advances in our understanding of the disease.

## Materials and Methods

### Fecal Sample Collection and Preparation

Fecal samples were sourced from an existing United States Department of Agriculture APHIS sample repository. White-tailed deer (*Odocoileus virginianus*) for this study were depopulated, farmed, naturally infected CWD positive and negative animals from the same herds that came from six different US states, coded and classified as Midwest, West, South, and East (Table S1a). Feces were collected manually from the rectum of each animal using a new nitrile glove to prevent cross-contamination. Feces was then placed into 50 mL conical tubes and stored at −80° C until aliquots were removed for this study.

The CWD status of the samples was determined by immunohistochemistry (IHC) of the medial retropharyngeal lymph nodes and obex of the brain at the USDA National Veterinary Services Laboratory in Ames, Iowa as previously described (*26*). An example of the diagnostics reults is shown in Fig. S1. CWD-positive samples were further subdivided into those with disease prions detected in both obex and lymph nodes (BRLN) and those with disease prions in lymph nodes but not in obex (LN), which were considered as possible cases of early and late stages of the disease. Those distinctions were included in some of the analysis as indicated in the text and figures.

### DNA Extraction and metagenomic sequencing

For DNA extraction, the automated KingFisher system was used with the MagMAX Microbiome Ultra Nucleic Acid Isolation Kit (ThermoFisher Scientific, A42356). Sample amount used for DNA extraction: 50 mg. Libraries prepared by targeted amplification of the variable V3-V4 regions of the bacterial 16S rRNA gene and attachment of Nextera XT indexes (Illumina Catalog #FC-131-2001) with PrimeStar Taq DNA Polymerase (Takara, cat#R045A). Concentrations were measured using the Qubit dsDNA HS (Invitrogen, Cat# Q32851). Samples were pooled, then purified using GenElute PCR Clean-Up Kit (Sigma, SKU NA1020), and KAPA Pure Beads (Roche-07983298001). Final library concentration determined by Invitrogen Qubit, and final size determined by Agilent DNA 1000 Kit (Agilent, #5067-1504) on the Agilent Bioanalyzer.

The gene-specific sequences used in this protocol target the 16S V3 and V4 regions. Sequencing primers were designed based on these gene-specific sequences with the addition of Illumina adapter overhang nucleotide sequences, to produce the following full-length primers: 16S Amplicon PCR Forward Primer: TCGTCGGCAGCGTCAGATGTGTATAAGAGACAGCCTACGGGNGGCWGCAG 16S Amplicon PCR Reverse Primer: GTCTCGTGGGCTCGGAGATGTGTATAAGAGACAGGACTACHVGGGTATCTAATCC The library pools were sequenced on Illumina® MiSeq™ with a V2 reagent kit, 500 cycles, with a Nano Flowcell (Illumina, MS-103-1003). Paired-end sequencing (2X250 cycles) was applied.

### Targeted metabolomics

Samples were lyophilized for 48 hours prior to metabolite extraction. Bile acids were extracted with 50% ACN spiked with the isotopically labeled standards cholic acid-*d*_5_, lithocholic acid-*d*_5_, sodium taurocholate-*d*_4_ and sodium taurodeoxycholate-*d*_4_ at concentrations of 2, 20, 1 and 1 µM, respectively. Bile acids were separated on a Waters ACQUITY UPLC BEH C_18_ column (2.1 mm × 150 mm, 1.7 μm, 130 Å) using a mobile phase of (A) 95% H2O 5% acetonitrile + 0.1% formic acid, and (B) 100% acetonitrile + 0.1% formic acid. The metabolites eluted with a linear gradient of 30% to 70% B over 16 min, at flow rate of 0.38 mL/min.

The short-chain fatty acids (SCFAs) were extracted with 70% IPA spiked with 50 µM of the isotopically labeled standards sodium acetate-^13^C_2_, sodium propionate-^13^C_3_ and sodium butyrate-^13^C_4_. The extracts were derivatized with 5 µL of 10 M pyridine, 10 µL of 250 mM *N*-(3-dimethylaminopropyl)-*N*’-ethylcarbodiimide (EDC) solution in 70% IPA and 10 µL of 250 mM 3-nitrophenylhydrazine (3-NPH) solution in 70% IPA. The mixture was reacted at 40 °C for 30 min. The reaction was quenched by adding 1.9 µL of formic acid. The SCFAs were separated with the same column and mobiles phases as the BAs, with the exception of a linear gradient of 10.0% to 47.5% B over 10.5 min, at flow rate of 0.38 mL/min.

The amino acids were extracted with water spiked with 50 µM of an isotopically labeled mix of 17 amino acids (Cambridge Isotope Laboratories). The amino acids were separated on a Imtakt Intrada Amino Acid column (3 mm × 150 mm, 3 μm) using a mobile phase of (A) acetonitrile + 0.3% formic acid, and (B) 20% acetonitrile 80% water + 80 mM ammonium formate. The amino acids eluted with a linear gradient of 20% B to 55% A over 12 min, at flow rate of 0.6 mL/min.

All of the LC-MS data was acquired on an Agilent 6460 Triple Quadrupole mass spectrometer with an electrospray ionization (ESI) source coupled with an Agilent 1290 Infinity II UPLC system. The LC-MS data was processed with Skyline v21.1 (MacCoss Lab, University of Washington).

### Bioinformatics Analysis

#### Read-level quality control

The 16S (V3-V4) paired-end amplicon sequences from 100 samples having read length of maximum 251 bases were quality assessed, followed by trimming and quality-based filtering using BBDuk (Bushnell, Brian. *BBMap: A Fast, Accurate, Splice-Aware Aligner. United States: N. p.,*2014; https://www.osti.gov/servlets/purl/1241166; http://jgi.doe.gov/data-and-tools/bb-tools/) to yield reads with average quality of Phred score(Q)>=30. The post trimming and filtering, quality report was assessed on reads level, base positions, length distribution etc.

### Taxonomic profiling

The quality trimmed fastq files were further processed using M-CAMP^TM^ web platform (*M-CAMP™: A cloud-based web platform with a novel approach for species-level classification of 16S rRNA microbiome sequences*. Schriefer et al. bioRxiv, 2021.08.25 456838; doi: https://doi.org/10.1101/2021.08.25.456838). The M-CAMP^TM^ uses the hybrid approach of heuristic alignment and k-mer based classification using kraken2 software (*27*) (https://ccb.jhu.edu/software/kraken2/). Reads were mapped against the proprietary “Sigma-Aldrich-16S_V3-V4” reference database which contains the V3-V4 primer specific 16S gene sequence. In the end, 76.02% to 89.91% raw reads in each sample were mapped to microbial sequences (Table S1b, Table S2). Following this analysis, the data were independently reanalyzed using QIIME2 (https://qiime2.org/), and the results of the two analyses were broadly compared. They are in agreement.

### Biodiversity analysis

For each sample, the different alpha-diversity indices of microbial species were calculated, including Pielou (for species evenness), Chao1 and observed features (for species richness), and Shannon and Simpson (for both species richness and species evenness) based on mapped taxa (rarefied to 36893, chosen to the maximum without losing samples, Table S3a); these indices and Faith’s phylogenetic diversity (PD) were also calculated based on the clustered reads and the aligned phylogenetic tree (rarefied to 30094, Table S3b). The significance for the difference between CWD-postive and CWD-negative alpha-diversity indices was tested with Wilcoxon rank sum test. The association between the disease state and the microbial composition was further studied using 2 beta-diversity measures: the Bray-Curtis dissimilarity matrix based on the mapped taxa (rarefied to 36893, Table S4a) and the variance-adjusted weight normalized Unifrac distance matrix based on the clustered reads and the aligned phylogenetic tree (rarefied to 30094, Table S4b), as well as the corresponding 2 matrices for a subset of samples (from the regions of Midwest 5, Midwest 7, and South 1) with higher rarefying factors (54026 for Table S4c and 47365 for Table S4d, respectively). PERMANOVA method of QIIME2 using 999 permutations was performed to assess the significance of pseudo-F statistics shown in Table 1. We further performed an analysis of variations due to combinations of different factors with R^2^ PERMANOVA analysis of QIIME2 using 999 permutations shown in Table 2 with the following fields of output: Df: Degree of freedom; SumsOfSqs: the sum of squares of deviation from the mean; F.Model: The F-statistic is a ratio of two variances; R^2^: statistical measure of the effect size (e.g., R^2^ of 0.25 means that 25% of the variation is explained by the grouping being tested); Pr(>F): p-value.

**Table 1.**
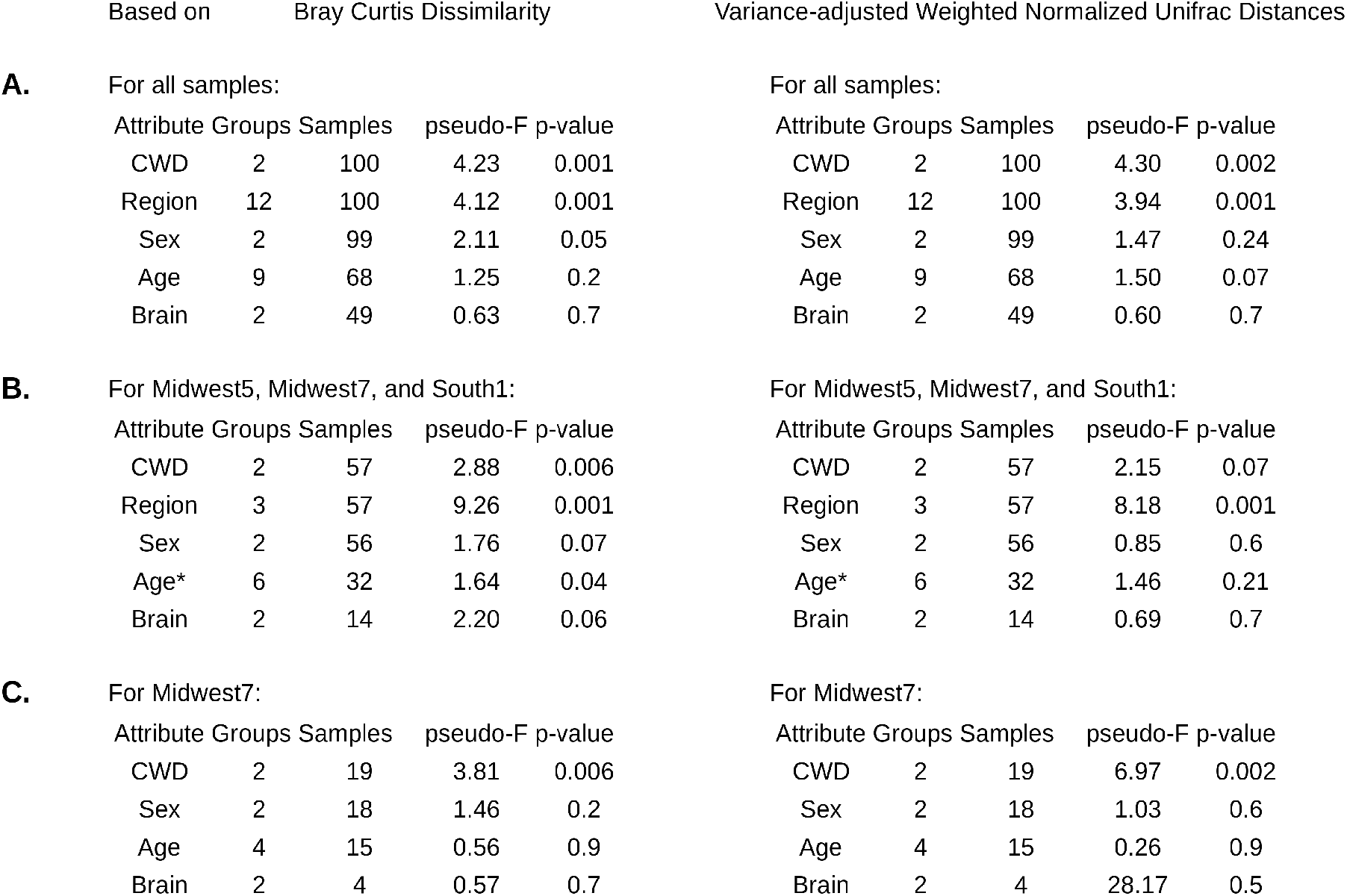
CWD status significantly affects beta-diversity of fecal microbiome. A. Beta-diversity changed significantly due to both CWD status and regions of origin of the samples. B. Beta-diversity also changed significantly due to regions of origin as in (A) for the samples from the 3 regions with both CWD-positive and CWD-negative deer. C. Beta-diversity changed significantly due to CWD status for the samples from one region. The statistics were calculated with PERMANOVA. See Table S4a-b for the matrices in A and Table S4c-d for in the matrices in B and C. * - age in South1 region was unknown.

**Table 2.** CWD status significantly affects beta-diversity of fecal microbiome. Analysis of variance of beta-diversity was performed for the 3 regions of Midwest5, Midwest7, and South1. A. Region of origin is still a significant factor after accounting for CWD status. B. CWD status is still a significant factor after accounting for region of origin. The statistics were calculated with PERMANOVA. See Table 4c-d for the the matrices.

| Based on |  | Bray Curtis Dissimilarity |  |  |  |  | Variance-adjusted Weighted Normalized Unifrac Distances |  |  |  |  |  |
| --- | --- | --- | --- | --- | --- | --- | --- | --- | --- | --- | --- | --- |
| A. |  | Df | SumsOfSqs | F.Model | R<sup>2</sup> | Pr(>F) |  | Df | SumsOfSqs | F.Model | R<sup>2</sup> | Pr(>F) |
|  | CWD | 1 | 0.38 | 3.81 | 0.050 | 0.004 | CWD | 1 | 0.086 | 2.93 | 0.038 | 0.018 |
|  | Region | 2 | 1.76 | 8.91 | 0.23 | 0.001 | Region | 2 | 0.52 | 8.93 | 0.23 | 0.001 |
|  | CWD:Region | 2 | 0.38 | 1.91 | 0.050 | 0.03 | CWD:Region | 2 | 0.18 | 3.11 | 0.080 | 0.003 |
|  | Residuals | 51 | 5.04 | NA | 0.67 | NA | Residuals | 51 | 1.49 | NA | 0.65 | NA |
|  | Total | 56 | 7.56 | NA | 1 | NA | Total | 56 | 2.29 | NA | 1 | NA |
| B. |  | Df | SumsOfSqs | F.Model | R<sup>2</sup> | Pr(>F) |  | Df | SumsOfSqs | F.Model | R<sup>2</sup> | Pr(>F) |
|  | Region | 2 | 1.93 | 9.76 | 0.26 | 0.001 | Region | 2 | 0.53 | 9.07 | 0.23 | 0.001 |
|  | CWD | 1 | 0.21 | 2.11 | 0.028 | 0.03 | CWD | 1 | 0.078 | 2.66 | 0.034 | 0.04 |
|  | Region:CWD | 2 | 0.38 | 1.91 | 0.050 | 0.04 | Region:CWD | 2 | 0.18 | 3.11 | 0.080 | 0.007 |
|  | Residuals | 51 | 5.04 | NA | 0.67 | NA | Residuals | 51 | 1.49 | NA | 0.65 | NA |
|  | Total | 56 | 7.56 | NA | 1 | NA | Total | 56 | 2.29 | NA | 1 | NA |

### Biomarker identification

LEfSe analysis was first performed to identify potential biomarkers (Table S5). Further statistical tests were performed to find the most prominent ones by filtering the list of detected taxa with abundance thresholds and evaluating the ones with the most significant p-values. For each taxa, p-values were calculated based on the relative abundance values (percentages) by the non-parametric Wilcoxon rank sum tests and, in geometric space (log), by two-tailed Welch’s t-test between CWD-positive and CWD-negative groups. For the ratio of each pair of taxa, p-values were calculated based on the ratio of their counts by Wilcoxon rank sum tests and, in geometric space (log), by two-tailed Welch’s t-test between CWD-positive and CWD-negative groups. Before calculating ratio and log, zero values were replaced with half of the non-zero minimum. The Benjamini-Hochberg false discovery rate (FDR) was calculated at each of the 7 rank level (Kingdom through Species). The calculated p-values and FDR are included in Table S6 and Table S8.

### Biomarker evaluation

We evaluated the separability of CWD-positive and CWD-negative samples using 2 techniques: the Linear Discrimination Analysis (LDA) (*42*) and the Support Vector Machine (SVC) (*43*). We found that LDA worked better than SVC for our purpose and we only present the LDA results in this paper. When separating samples into 2 groups, LDA is the same as Fisher linear discrimination analysis (*44*). Briefly, based on each set of biomarkers, we performed the separation/identification of CWD-positive and CWD-negative samples by thresholding a weighted projection of the sample values of the markers (e.g. abundance values of taxa) in geometric space (log). The weights were proportional to the Fisher Linear Discriminant (FLD) and the threshold was half way between the two projected averages of the CWD-positive and CWD-negative training sets. To evaluate the performance on test data, we used jackknife (leave one out) re-sampling technique to calculate the identification rates. Unless specifically stated otherwise, we only present the identification rates on test data, which are different from, normally lower than, the rates on training data. To visualize the data, we also calculated the second FLD in the subspace orthogonal to the first and etc. This was programmed in Matlab (version R2023b).

### Cross-omics analysis

The bacterial relative abundances were filtered to include enriched bacteria only. The metabolite amounts (normalized to isotopically labelled standards) were correlated to the filtered bacteria relative abundances using Spearman’s rank correlation. The heatmap was generated using the pheatmap R package (version 1.0.12).

### Data availability

The 16S rRNA sequencing data is available in the NCBI Sequence Read Archive with the BioProject ID: PRJNA936583.

## Results

### Analysis of deer feces microbial composition

To identify potential changes in the microbial composition of the fecal samples in response to CWD, we isolated total DNA from fecal pellets of 50 CWD-negative and 50 CWD-positive depopulated farmed white-tailed deer. CWD-positive (pos) deer in this set were diagnosed by immunohistochemistry of paraffin-embedded sections of brain (BR) and/or lymph node (LN), and the disease status was classified as CWD-negative (neg) for those deer that showed no prion accumulation in either of these tissues. An example of brain diagnostics is shown in Fig. S1.

DNA extracted from individual fecal samples was amplified using targeted amplification of the variable V3-V4 regions of the bacterial 16S rRNA gene, and sequenced on the Illumina platform to identify their microbial composition. After stringent filtering for clean reads and discarding the data below 0.01% abundance in the sample, 892 different microbial taxa were present across the analyzed samples, shown in the comprehensive and specific rank-wise (Level 1: Kingdom to Level 7: Species) taxon count table summarizing the count for each observed taxon (Table S2a) and the abundance table of the clade percentage, i.e., clade fragment (the accumulative or collapsed amount of taxa in each sample at different taxonomic levels) percentage of the total in each sample (Table S2b).

We first analyzed alpha-diversity, i.e. the species diversity of microbial communities in each individual sample. This analysis revealed no statistically significant difference in alpha-diversity between CWD-positive and CWD-negative samples based on the mapped taxa (Table S3a) and based on the clustered reads and the aligned phylogenetic tree (Fig. S2, Table S3b).

### Beta-diversity of microbial composition of the deer feces is highly affected by CWD

We next analyzed beta-diversity, i.e. the species diversity between microbial communities in different samples, measured by Bray-Curtis dissimilarity matrix based on observed taxa as well as variance-adjusted weight normalized Unifrac distance matrix based on clustered reads and phylogenetic tree. We performed principal coordinate analysis (PCoA) and t-distributed stochastic neighbor embedding (tSNE) to visualize the beta-diversity (Fig. 1A and 1B, respectively) and determined the significance of the dissimilarity and the distance between CWD-positive and CWD-negative samples with PERMANOVA statistical test. We found that the disease status significantly affect the beta-diversity (Table 1A).

**Figure 1.**
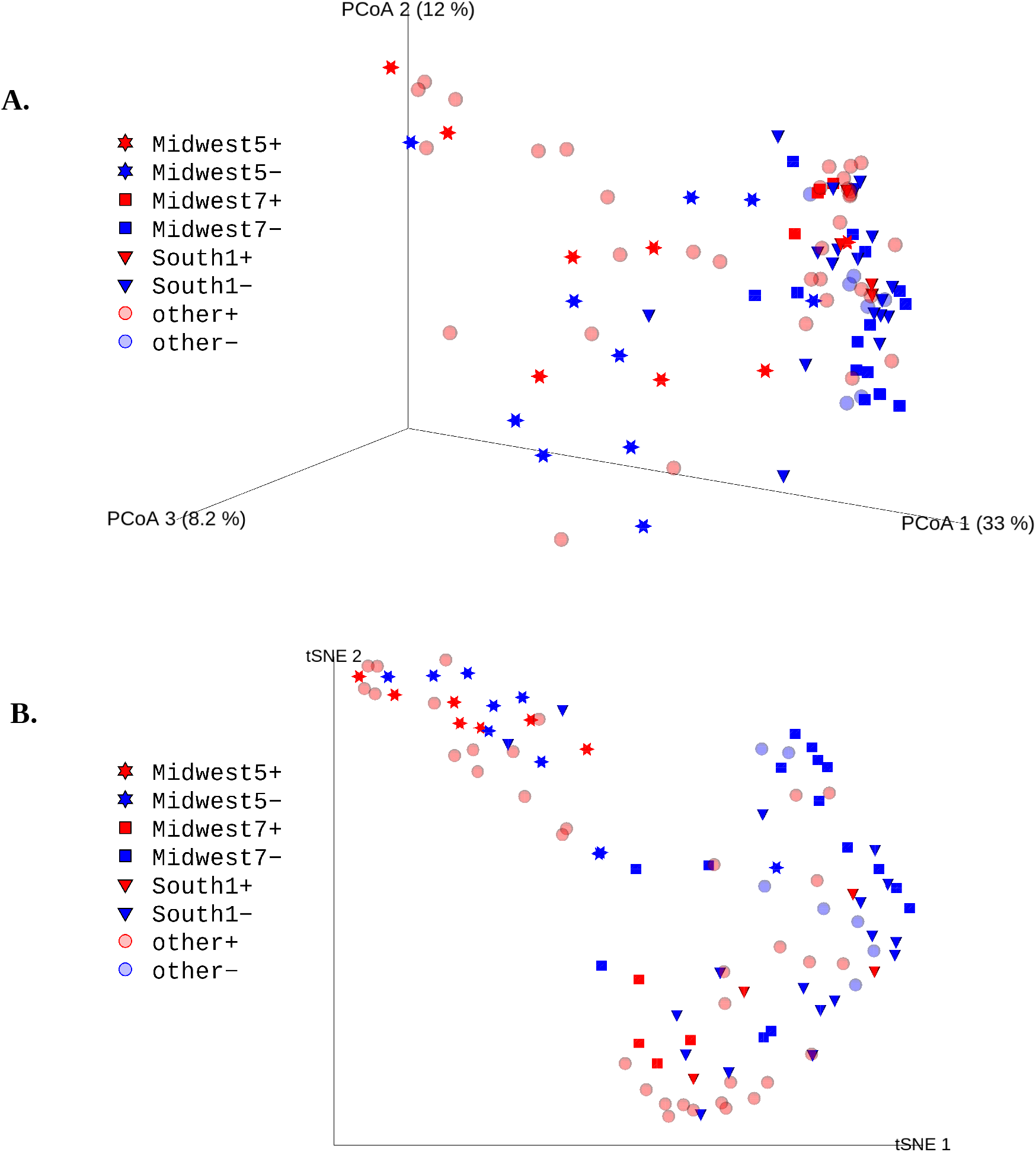
CWD status significantly affects fecal microbiome. A. and B. PCoA and tSNE plots of Bray-Curtis dissimilarity matrix of microbiomic composition of the deer fecal sample. The analysis was performed for the entire set of samples; the regions that had representation of both CWD-positive (+) and CWD-negative (-) samples (Midwest5, Midwest7, and South1) are marked by different symbols as indicated. Red, CWD positive. Blue, CWD negative.

Fecal samples used in this study come from deer different from each other by several other parameters, including sex, age, geographic region of origin, and presence of misfolded prions in the brain in addition to the lymph nodes (Table S1a). All of these parameters can potentially affect microbial composition. In particular, geographical origin is expected to strongly affect the repertoire of microorganisms found in fecal samples (35), given potentially large variability in microbial and plant ecology in different regions of the US. The PERMANOVA statistical test showed that geographic origin of the deer was indeed another significant factor driving beta-diversity between the samples (Table 1A), also illustrated in Fig. 1. In comparison, sex showed a border-line significance while age and disease prion in brain showed no significant effect (Table 1A).

Analysis of the sample metadata (Table S1a) indicated that the disease status and the region of origin could be potentially confounded by the fact that deer from some regions were exclusively CWD-positive, while in others deer were exclusively CWD-negative. Re-analysis of the samples limited to the 3 regions where both CWD-positive and CWD-negative samples were represented (Midwest5, Midwest7, and South1) showed that both the disease status and the region of origin still emerged as the 2 main factors driving a statistically significant beta-diversity between samples (Table 1B). Analysis of variance (Table 2) showed that geographical origin and CWD status both are the likely drivers for the differences in the microbial composition in fecal samples. Notably, CWD status shows promising trends suggesting that deer from the same region, analyzed in sufficient numbers, would show differences in their gut microbiomes that are driven by the disease (e.g., Table 1C and Fig. S3).

### Microbial composition of the deer feces exhibits distinct CWD-dependent microbial signatures

To identify specific microbial taxa that are differentially abundant between CWD positive and negative deer, we first performed LEfSe analysis over the 892 taxa. This analysis yielded a list of 90 differentially abundant taxa (Fig. S4 and Table S5a). To narrow the list down, we used more stringent methods for microbiome differential abundance (*45*). We filtered the 892 taxa down to 134 taxa which have the average relative abundance >0.1% in CWD-positive and/or CWD-negative samples. We then calculated the p-values according to the clade percentage.

Since multiple statistical tests were performed and the taxon counts were used repeatedly at each rank level, we evaluated the false discovery rate (FDR) at each of the 7 rank levels. These methods of analysis yielded 29 and 27 taxa with p-value <0.05 and FDR <0.1 for Welch’s t-test and Wilcoxon rank-sum test, respectively, and with 25 in common (Table S6). They are also in common with the LEfSe analysis, 25 and 22, respectively.

This analysis revealed several interesting trends. First, at each taxonomic level, microbial taxa grouped into distinct abundance patterns, with some taxa lower and others higher in either CWD-positive or CWD-negative samples (Fig. 2, Table 3). Second, these patterns could potentially be used to identify the disease status; for example, the 6 differentially abundant taxa at the Genus level comprised ∼4% of the observed microbiota (Fig. 2, 3A) and could be used to discriminate between CWD-positive and CWD-negative samples (Fig. 3B-C)---the first Fisher Linear Discriminant (FLD) yielded identification rates of 74% and 73% for CWD-negative and CWD-positive deer. Third, many differentially abundant taxa conveyed consistent information at different rank levels as well as for individual region and sex; for example, Coriobacteriia Class with Coriobacteriales, Atopobiaceae, and Olsenella (Fig. 4A-D), and Clostridia Class with Clostridales, Eubacteriacease, and Dorea (Fig S5A-D) were consistently less abundant in CWD-positive samples at the rank levels of Class, Order, Family, and Genus; in addition, the FLD yielded similar identification rates for CWD-negative and CWD-positive samples at those 4 rank levels (Table 4A). Fourth, among CWD-positive samples which include 24 with disease prions in both obex and lymph nodes (BRLN) and 25 with disease prions in lymph nodes but not in obex (LN), we found that Coriobacteriia and Clostridia were consistently less abundant in both LN and BRLN samples (Fig. 4E-F and Fig. S5E-F, respectively) and the FLD worked similarly for LN and BRLN samples (Table 4A).

**Figure 2.**
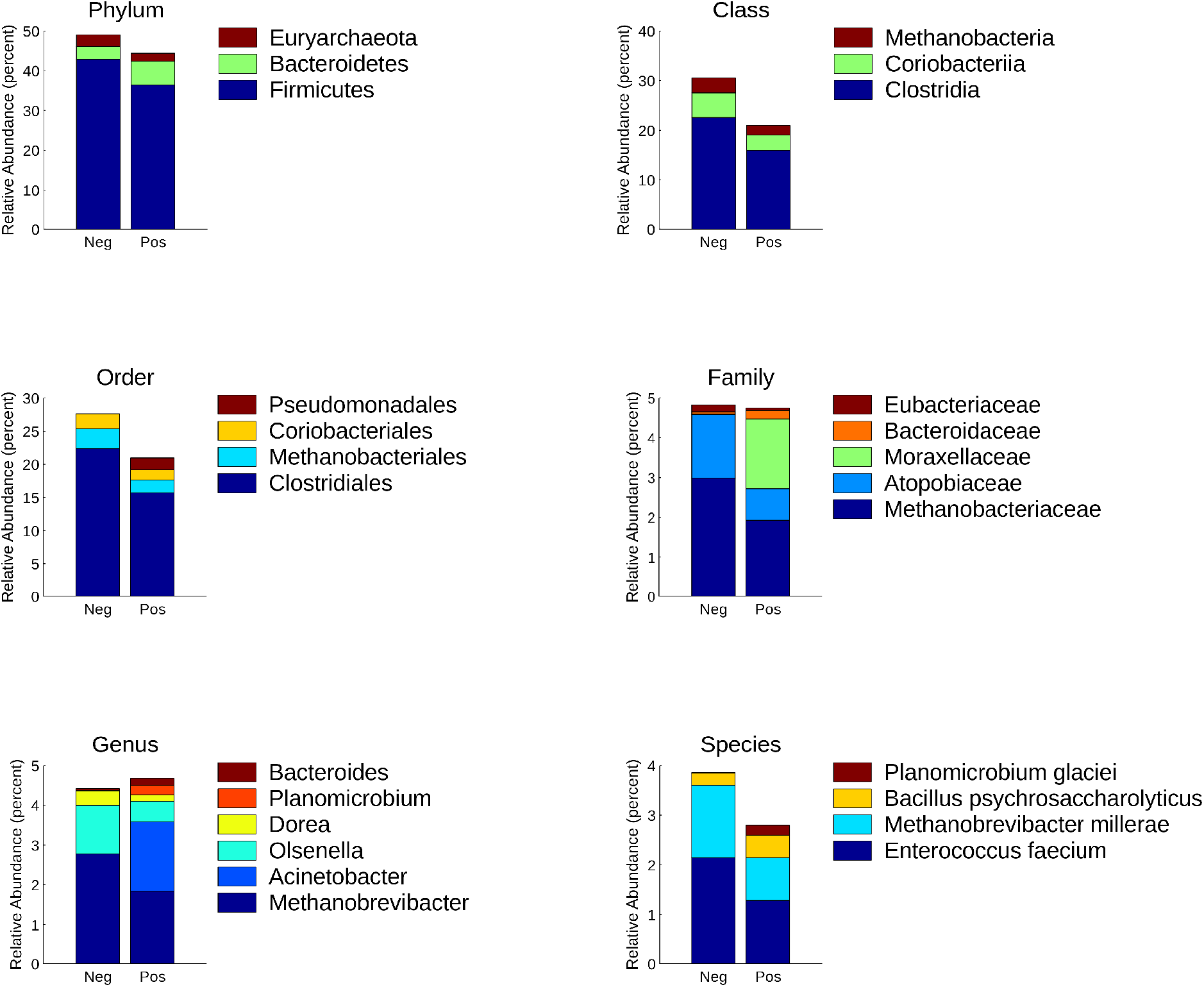
Differences in the microbiota between feces from CWD-positive and CWD-negative deer at different taxonomic levels. Significantly changed taxa (Wilcoxon rank sum test p<0.05 and FDR<0.1) with >0.1 average abundance (%) in at least one type of the fecal samples are shown. Bar charts show the average abundance of each taxa in the CWD-negative (Neg) versus CWD-positive (Pos) samples. See Table 3 for details. See Table S6 for the list of all taxa with >0.1 average abundance.

**Figure 3.**
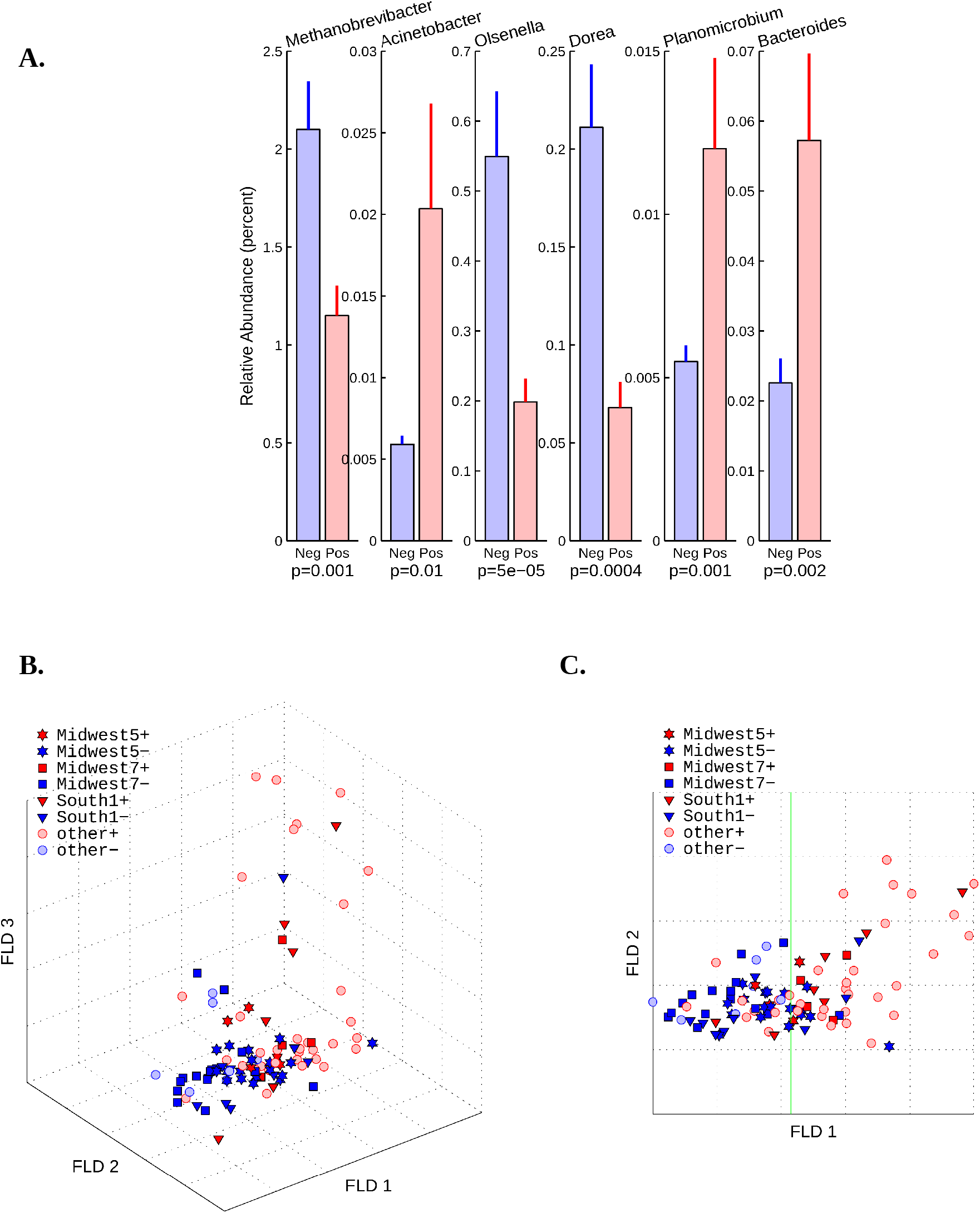
Taxa as potential biomarkers of CWD. A. The bar charts show the average abundances of taxa in Genus with a significant CWD-dependent abundance. Bars represent geometric means. Error bars represent SEM, n=50 for each set. The p-values are from Wilcoxon rank sum test. B. 3-D Fisher Linear Discriminant (FLD) projection of those abundances shows the clustering of CWD-positive (red) and CWD-negative (blue) samples. C. 2-D projection with the green line indicating the threshold for FLD 1.

**Figure 4.**
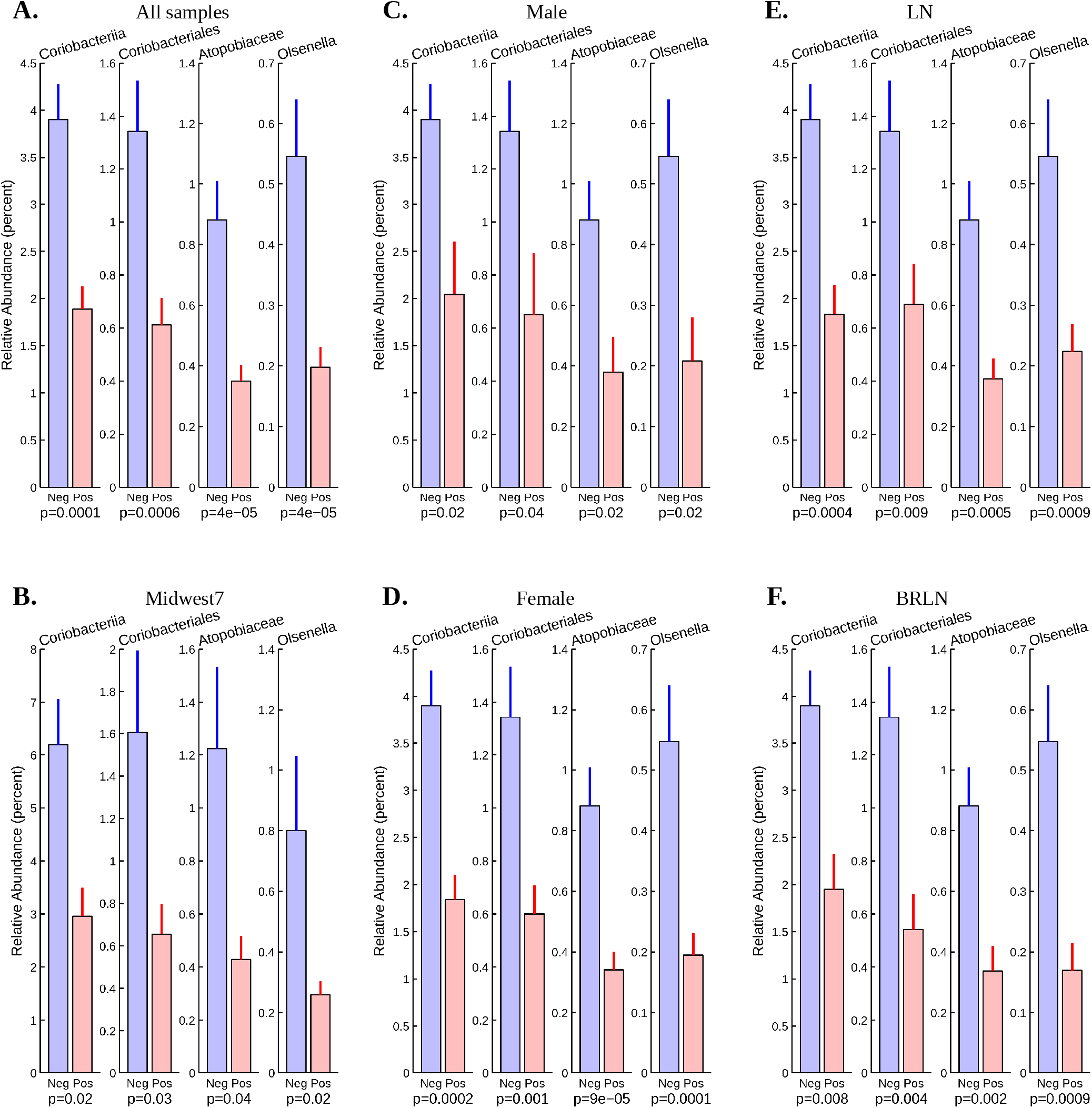
Coriobacteriia are consistently less abundant in CWD-positive samples. CWD-positive samples show a significant reduction in abundance of the Coriobacteriia Class, the Coriobacteriales Order, the Atopobiaceae Family, and the Olsenella Genus. A. In all samples. B. In samples from Midwest7 region with both CWD-positive and CWD-negative deer. C. In male samples. D. In female samples. E. Positive samples with mis-folded prions in LN only. F. Positive samples with mis-folded prions in BR and LN. Bars represent geometric means. Error bars represent SEM. The p-values are from Wilcoxon rank sum test.

**Table 3.** 27 abundant and significantly changed taxa due to CWD disease. Significantly changed taxa (Wilcoxon rank sum test p<0.05 and FDR<0.1) with >0.1% average abundance in at least one type of the fecal samples are shown. K_: Kingdom. p_: Phylum. c_: Class. o_: Order. f_: Family. g_: Genus. s_: Species. Mean_neg: average abundance of CWD-negative. Mean_pos: average abundance of CWD-positive. See Table S6 for the list of all taxa with >0.1% abundance values.

| Scientific Name | NCBI id | P value | FDR | Mean_neg | Mean_pos |
| --- | --- | --- | --- | --- | --- |
| k__Archaea | 2157 | 0.0013 | 0.0038 | 3 | 1.9 |
| k__Bacteria | 2 | 0.0064 | 0.0096 | 96 | 97 |
| p__Bacteroidetes | 976 | 0.025 | 0.099 | 3.2 | 6.1 |
| p__Euryarchaeota | 28890 | 0.0013 | 0.015 | 3 | 1.9 |
| p__Firmicutes | 1239 | 0.0029 | 0.018 | 43 | 36 |
| c__Clostridia | 186801 | 0.00097 | 0.006 | 23 | 16 |
| c__Coriobacteriia | 84998 | 0.00011 | 0.0011 | 5 | 3 |
| c__Methanobacteria | 183925 | 0.0012 | 0.006 | 3 | 1.9 |
| o__Clostridiales | 186802 | 0.00078 | 0.0054 | 22 | 16 |
| o__Coriobacteriales | 84999 | 0.00055 | 0.005 | 2.3 | 1.5 |
| o__Methanobacteriales | 2158 | 0.0012 | 0.0068 | 3 | 1.9 |
| o__Pseudomonadales | 72274 | 0.018 | 0.065 | 0.0084 | 1.8 |
| f__Atopobiaceae | 1643824 | 3.9E-05 | 0.00043 | 1.6 | 0.79 |
| f__Bacteroidaceae | 815 | 0.00021 | 0.0015 | 0.056 | 0.21 |
| f__Eubacteriaceae | 186806 | 8.4E-05 | 0.00074 | 0.17 | 0.061 |
| f__Methanobacteriaceae | 2159 | 0.0012 | 0.0067 | 3 | 1.9 |
| f__Moraxellaceae | 468 | 0.0065 | 0.031 | 0.0083 | 1.8 |
| g__Acinetobacter | 469 | 0.012 | 0.056 | 0.0083 | 1.7 |
| g__Bacteroides | 816 | 0.0019 | 0.011 | 0.044 | 0.18 |
| g__Dorea | 189330 | 0.00036 | 0.0032 | 0.37 | 0.17 |
| g__Methanobrevibacter | 2172 | 0.0011 | 0.0081 | 2.8 | 1.8 |
| g__Olsenella | 133925 | 4.5E-05 | 0.00071 | 1.2 | 0.52 |
| g__Planomicrobium | 162291 | 0.0013 | 0.0098 | 0.012 | 0.23 |
| s__Bacillus psychrosaccharolyticus | 1407 | 0.012 | 0.063 | 0.25 | 0.46 |
| s__Enterococcus faecium | 1352 | 0.0094 | 0.045 | 2.1 | 1.3 |
| s__Methanobrevibacter millerae | 230361 | 0.00064 | 0.0073 | 1.5 | 0.86 |
| s__Planomicrobium glaciei | 459472 | 0.0013 | 0.012 | 0.011 | 0.2 |

**Table 4.** FLD classifiers at the rank levels correctly identify CWD in the majority of the test samples. Each X/Y entry in the tables is the ratio of the number of correctly identified ones over the test of 50 CWD-negative (CWD-neg) samples and 49 CWD-positve (CWD-pos) samples (consisted of 25 LN and 24 BRLN). When training an FLD classifier, all samples were used except one which was left to test if the FLD correctly classified it as CWD-positive or CWD-negative. The taxa used for identification at each of the rank levels in A, B, C and D are shown in Table 3, Table S5a, and Table S6. For E, the indicated number of taxa ratios at each rank level with the smallest Welch’s t-test p-values was used (Table S8).

| **A. Taxa** |  |  |  |  |  |
| --- | --- | --- | --- | --- | --- |
| Rank | # of Taxa | CWD-neg | CWD-pos | LN | BRLN |
| Phylum | 3 | 35/50 | 32/49 | 18/25 | 14/24 |
| Class | 3 | 35/50 | 33/49 | 18/25 | 15/24 |
| Order | 4 | 36/50 | 28/49 | 14/25 | 14/24 |
| Family | 5 | 37/50 | 33/49 | 17/25 | 16/24 |
| Genus | 6 | 37/50 | 36/49 | 21/25 | 15/24 |
| Species | 4 | 38/50 | 34/49 | 16/25 | 18/24 |

| **B. LEfSe** |  |  |  |  |  |
| --- | --- | --- | --- | --- | --- |
| Rank | # of Taxa | CWD-neg | CWD-pos | LN | BRLN |
| Phylum | 5 | 38/50 | 27/49 | 16/25 | 11/24 |
| Class | 6 | 36/50 | 30/49 | 16/25 | 14/24 |
| Order | 11 | 31/50 | 33/49 | 16/25 | 17/24 |
| Family | 15 | 35/50 | 35/49 | 17/25 | 18/24 |
| Genus | 29 | 37/50 | 32/49 | 16/25 | 16/24 |
| Species | 21 | 39/50 | 35/49 | 17/25 | 18/24 |

| **C. Abundance>0.1** |  |  |  |  |  |
| --- | --- | --- | --- | --- | --- |
| Rank | # of Taxa | CWD-neg | CWD-pos | LN | BRLN |
| Phylum | 10 | 35/50 | 30/49 | 17/25 | 13/24 |
| Class | 13 | 30/50 | 34/49 | 17/25 | 17/24 |
| Order | 18 | 33/50 | 33/49 | 19/25 | 14/24 |
| Family | 28 | 39/50 | 34/49 | 17/25 | 17/24 |
| Genus | 36 | 37/50 | 35/49 | 18/25 | 17/24 |
| Species | 27 | 37/50 | 31/49 | 16/25 | 15/24 |

| **D. Abundance>1.0** |  |  |  |  |  |
| --- | --- | --- | --- | --- | --- |
| Rank | # of Taxa | CWD-neg | CWD-pos | LN | BRLN |
| Phylum | 7 | 35/50 | 28/49 | 17/25 | 11/24 |
| Class | 10 | 30/50 | 32/49 | 18/25 | 14/24 |
| Order | 13 | 37/50 | 34/49 | 20/25 | 14/24 |
| Family | 15 | 35/50 | 33/49 | 15/25 | 18/24 |
| Genus | 12 | 37/50 | 33/49 | 16/25 | 17/24 |
| Species | 8 | 38/50 | 32/49 | 16/25 | 16/24 |

**Table 4. FLD classifiers at the rank levels correctly identify CWD in the majority of the test samples.**
| **E. Taxa Ratio** |  |  |  |  |  |  |
| --- | --- | --- | --- | --- | --- | --- |
| Rank | # of Taxa | CWD-neg | CWD-pos | LN | BRLN | # of Ratio |
| Phylum | 5 | 32/50 | 28/49 | 17/25 | 11/24 | 4 |
| Class | 6 | 33/50 | 31/49 | 18/25 | 13/24 | 5 |
| Order | 5 | 41/50 | 24/49 | 12/25 | 12/24 | 4 |
| Family | 6 | 40/50 | 31/49 | 15/25 | 16/24 | 5 |
| Genus | 6 | 39/50 | 32/49 | 16/25 | 16/24 | 5 |
| Species | 5 | 36/50 | 32/49 | 15/25 | 17/24 | 5 |

To fully evaluate these taxa as suitable biomarkers, we used FLD to calculate the identification rates at each rank level and found that these taxa achieved rates ∼70%. It is important to emphasise that even though the number of these identified markers at each rank level (Table 4A) is a fraction of the ones identified by LEfSe (Table 4B) or by the abundance>0.1 filter (Table 4C), they still achieved the robust, promising performance on test data on par with LEfSe and the abundance>0.1 filter.

### Metabolic composition of the deer feces exhibits significant CWD-dependent changes

To see if the changes in microbiota are accompanied by changes in metabolite composition in the feces, we performed targeted metabolomics to detect short-chain fatty acids, amino acids, and bile acids in deer fecal samples. This analysis identified 54 metabolites (12, 31, and 11, respectively, see Table S7a) in the samples. Some of these metabolites showed significant differences between feces of CWD-positive and CWD-negative deer (Fig. 5A) and could be used to discriminate the samples according to the disease status (Fig. 5B), with the first Fisher Linear Discriminant (FLD) yielding identification rates of 92% and 77% for CWD-negative and CWD-positive deer, respectively.

**Figure 5.**
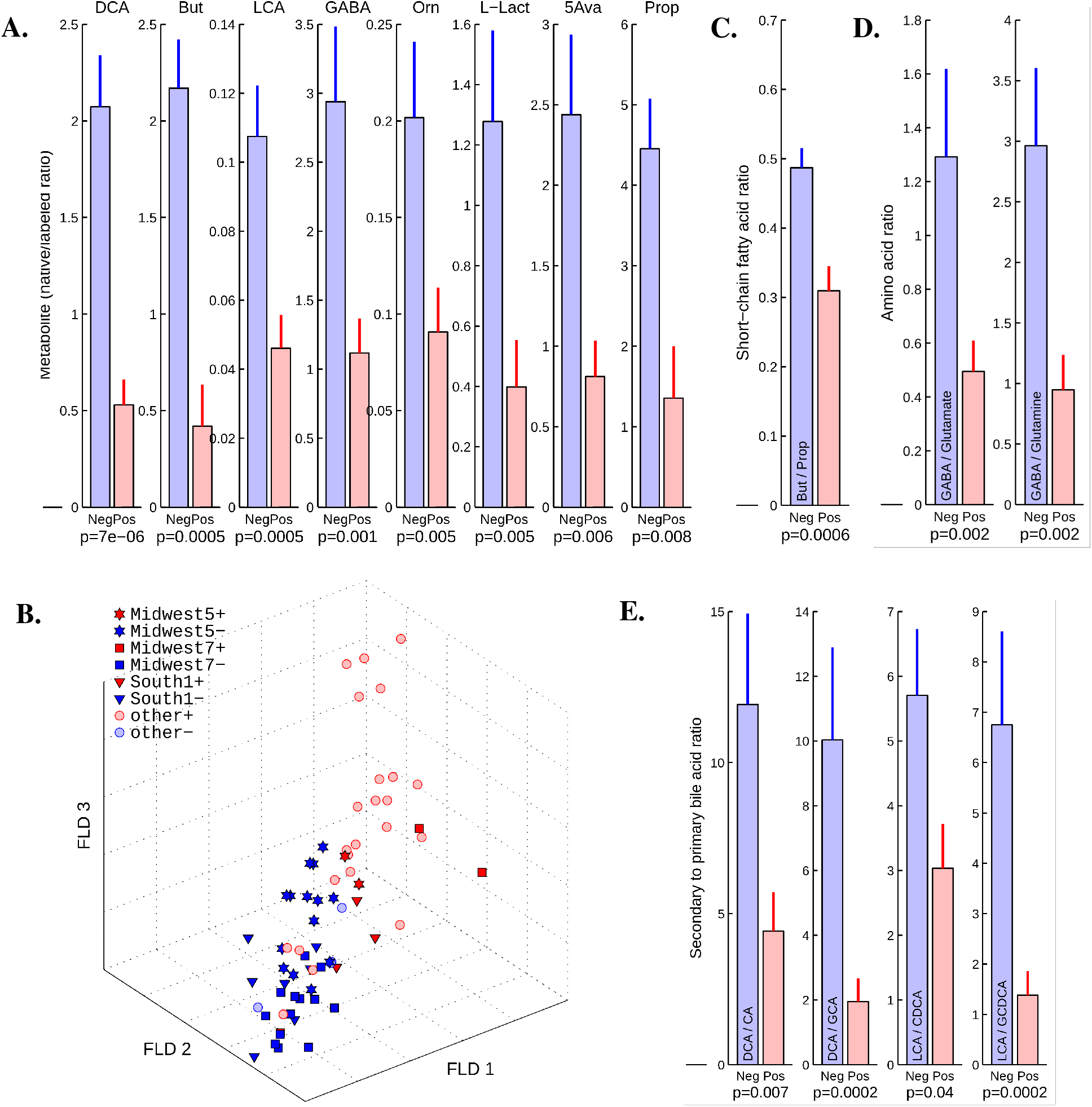
Difference between metabolites in feces of CWD positive and negative deer. A. Specific metabolites show significant CWD-dependent changes (Wilcoxon rank sum test p<0.01 and FDR<0.1). B. 3D Fisher Linear Discriminant (FLD) projection. C. Butyric to propionic acids ratio. D. GABA to glutamate and glutamine ratios. E. Secondary to primary bile acid ratios. The bars represent the geometric means of the indicated metabolites in A, and their ratios in C-E in CWD-negative (Neg, n=38) versus CWD-positive (Pos, n=31) samples. Error bars represent SEM. 5Ava - 5-Aminopentanoic acid, But – butyric acid, CA - cholic acid, CDCA - glycodeoxycholic acid, DCA - deoxycholic acid, GABA - gamma-aminobutyric acid, GCA - clycocholic acid, GCDCA - glycochenodeoxycholic acid, LCA - lithocholic acid, L-Lact - l-lactic acid, Orn – ornithine, Prop – propionic acid.

Next, we quantified relative amounts of some of these metabolites in CWD-positive versus CWD-negative feces. The ratio of butyric to propionic short-chain fatty acid (Fig. 5C) and the ratios of amino acid GABA to the excitatory neurotransmitter glutamate and its precursor glutamine (Fig. 5D) were significantly decreased in CWD-positive feces. The secondary bile acid DCA to the primary bile acids CA and GCA ratios as well as LCA to CDCA and GCDCA ratios were also significantly decreased (Fig. 5E). These changes in the ratios of metabolite levels also have a potential use as biomarkers for CWD diagnostics. Using the 7 ratios in Fig. 5C-E, the FLD yielded identification rates of 89% and 74% for CWD-negative and CWD-positive deer, respectively, similar to the rates from using the 8 metabolites in 5A.

For this metabolite analysis, we used 69 of the original 100 samples used for the microbiomics described in previous sections, chosen based on the sample abundance. Using the 69 samples, we performed cross-omics analysis between the microbial taxa and the identified metabolites. The cross-omic analysis (Fig. 6) shows correlations between 63 significantly enriched microbial taxa (Table S7c) and 54 metabolites (Table S7a). Among those, e.g., the relative abundance of the *Clostridia* and Coriobacteriia class had significant positive correlations (FDR <0.05) with the secondary bile acids, DCA and LCA (both less abundant in CWD-positive deer). These correlations are in agreement with the literature as these species are known to facilitate the conversion of primary bile acids to secondary bile acids (*30*).

**Figure 6.**
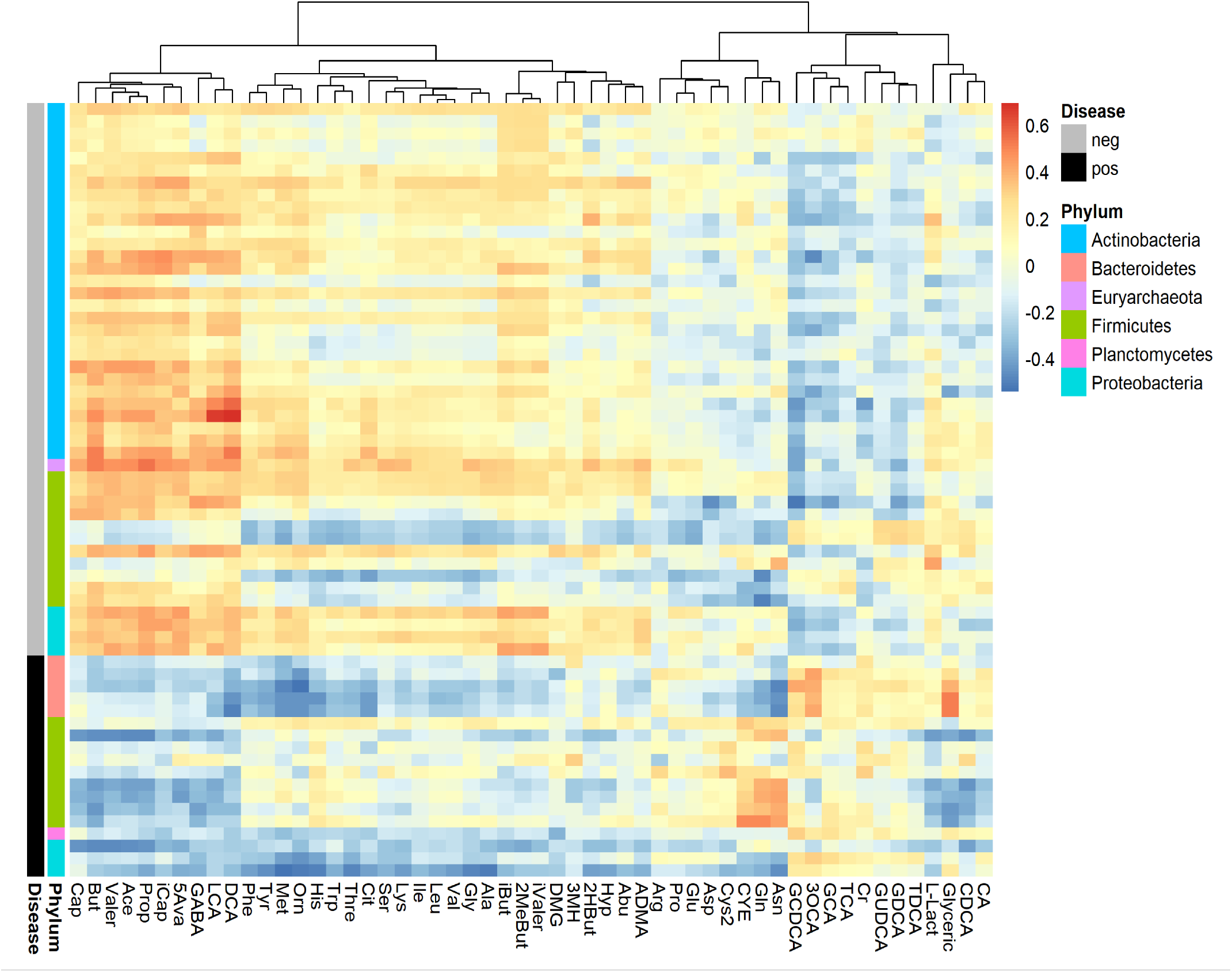
Heatmap of the correlation coefficient between metabolites and microbial abundance. The correlation coefficient of 63 significantly enriched microbial taxa (rows), to 54 metabolites (columns). The color-scaled heatmap corresponds to Spearman’s rank correlation coefficients. Only LEfSe enriched bacteria with significant correlations to metabolites (FDR<0.05) are shown. The microbial taxa (rows) are sorted for enriched in CWD-negative or CWD-positive fecal samples. Microbial taxa annotation represents the taxonomic profile at the Phylum level.

### Taxa ratios as potential diagnostic markers of CWD

The differences revealed by the individual taxa suggest that the microbiome changed in CWD. However, those analysis depends on the normalized expression values, i.e., the percentage of an expressed taxa. It would be more efficient to exploit the differences without measuring the entire microbiome of each sample.

Previous studies found that in some cases ratios of specific pairs of microbial taxa in each sample can be indicative of certain disease or physiological status. For example, Firmicutes to Bacteriodetes (F/B) ratio was previously proposed as a marker of obesity and type 2 diabetes (*31*), inflammatory bowel disease (*32*), and aging (*33*). In humans, higher F/B ratios (>12) are found in healthy individuals, suggesting that reduced F/B ratio may be a potential clinical biomarker (*34*). In deer feces, this ratio shows a similar trend: it is decreased in CWD positive animals, from 21 in CWD-negative to 12 in CWD-positive (one-tailed Welch’s t-test p=0.04, Wilcoxon rank sum test p=0.01). Clearly, more studies are needed to evaluate F/B ratio as potential diagnostic markers for CWD. However, the general idea of identifying ratios of specific pairs of taxa in CWD-positive versus CWD-negative samples, may be a promising approach that could potentially serve as a basis for a more rapid CWD scoring test compared to the full microbiomics signatures.

To test the potential of using ratios of the most abundant taxa in the fecal samples as CWD biomarkers, we filtered the 892 taxa further down to 67 taxa which have the average relative abundance >1% in CWD-positive and/or CWD-negative. It turned out that those taxa still contained the information to identify CWD disease with the rates (Table 4D) similar to the ones from LEfSe (Table 4C). We calculated the ratios for those taxa pairs within each rank, and found 65 ratios of taxa pairs with significantly higher ratios of CWD-negative to CWD-positive (fold-change>1.5, p-value<0.05, and FDR<0.1, Figs S6-S10 and Table S8). These ratios represent potential CWD-dependent signatures in the fecal microbiome that could be explored for CWD surveillance and diagnostics. Using the top 5 ratios (involving 6 taxa) at the Genus level (Fig. 7), the FLD yielded identification rates of 78% and 71% for CWD-negative and CWD-positive deer, respectively, similar to the rates (74% and 71%) from using 28 taxa identified by LEfSe at the Genus level.

**Figure 7.**
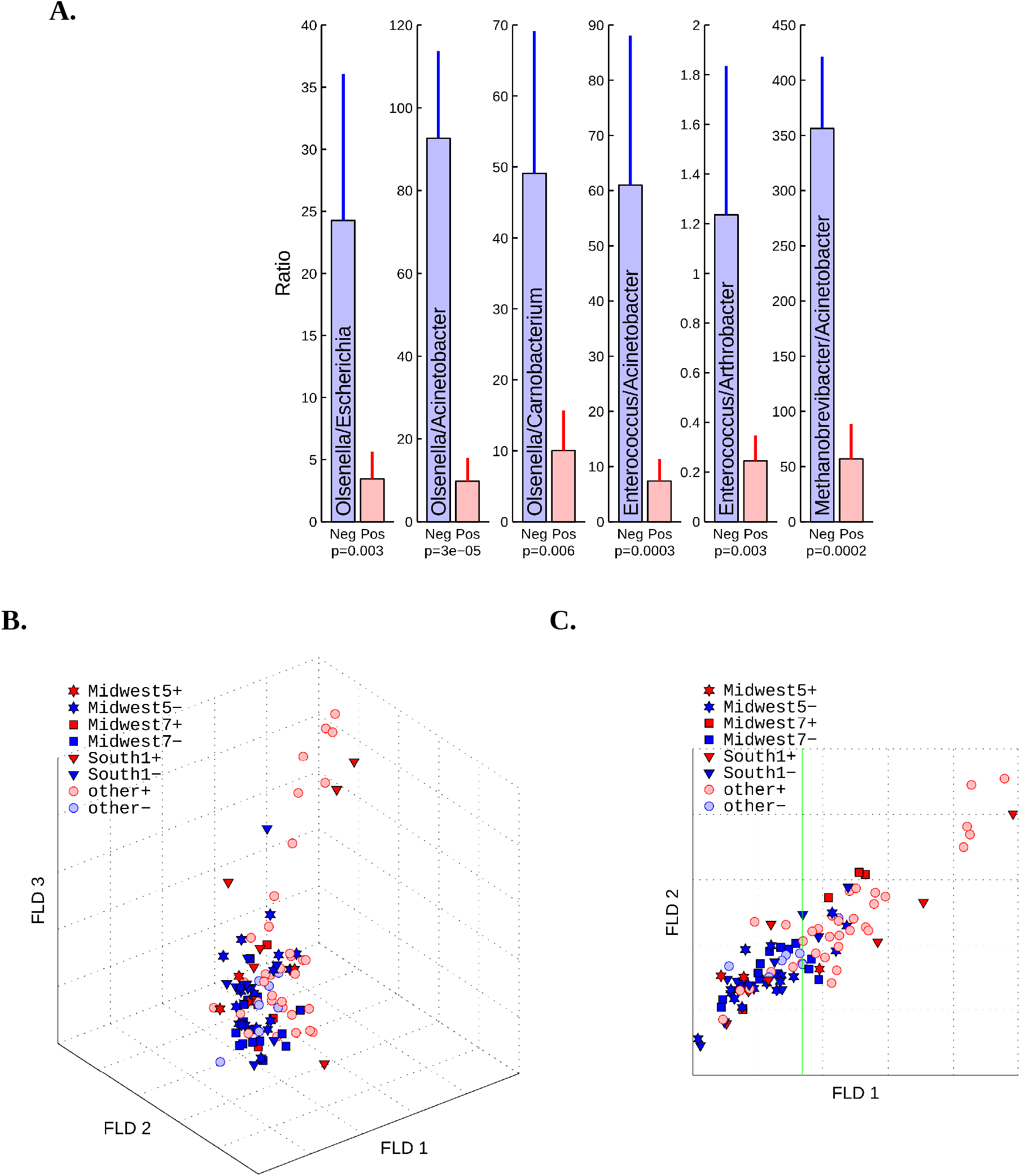
Taxa ratios as potential biomarkers of CWD. A. The bar charts show the geometric mean of the ratio of the indicated taxa pair of the top 6 at the Genus rank level with the most significant change in their ratio between CWD-negative (Neg) and CWD-positive (Pos) samples. Error bars represent SEM, n=50 for each set. The p-values are from two-tailed Welch’s t-test. B. 3-D Fisher Linear Discriminant (FLD) projection of those ratios shows the clustering of CWD-positive (red) and CWD-negative (blue) samples. C. 2-D projection with the green line indicating the threshold for FLD 1.

The full comparison (Table 4) shows that using taxa ratios as CWD biomarkers has the advantage of achieving the same identification performance with a less number of microbes to measure. This is especially true at the Species and Genus levels.

## Discussion

Our study represents a comprehensive analysis of fecal microbiome from deer with and without CWD, with the goal of identifying potential biomarkers that could be utilized for novel types of ante mortem CWD surveillance and diagnostics. We identified a total of 27 highly abundant microbial taxa that are differentially abundant between CWD-positive and CWD-negative feces (Table 3) and can be used as biomarkers for CWD identification at different rank levels with the the abundance values of a few taxa at each level (Table 4A). Furthermore, we found a correlation of these taxa changes with the changes in key metabolites in the feces. We also identified microbial taxa ratios that show promising trends in the future development of methods for CWD diagnostics using a few ratios at each rank level (Table 4E). While the variability of microbial taxa in the deer feces is also strongly influenced by geographical region, as well as likely by diet, seasonal changes, and other variables not included in our study, our data proposes a use of abundant microbial taxa as a tool to detect “microbiomic signatures” of CWD. This approach could eventually lead to breakthroughs in our understanding and control of CWD.

A recent study that performed a similar type of analysis using feces from ∼200 farmed deer, identified no robust changes in fecal microbiota associated with CWD, but did find differences related to the geographic origin of the deer, likely related to diet and partially correlated with sex (*35*). In our study, the geographical origin is also a major factor that drives fecal microbiome variability between the samples, e.g., Osenella and Dorea were identified as CWD markers in Midwest7 region, they were not in Midwest5 and South1 (Fig. 4B, Fig. S5B, and Table S5b-c). This is not surprising, given that environmental microbial composition differs greatly in different habitats. However, encouragingly, our data also points to potential patterns of changes in fecal microbiomes that may be directly linked to CWD. Evaluating these results in a larger sample set obtained from the same geographical region is essential for the identification of the global trends of the CWD-specific changes in fecal microbiomes.

The samples used in our study included those with disease prions in both obex and lymph nodes (BRLN) as well as those with disease prions in lymph nodes but not in obex (LN) (Table S1a). Since the disease prion likely propagates from the lymph nodes to the brain, the LN-positive deer likely have earlier stages of the disease compared to BRLN. Notably, the disease-related changes seen in our study were similar in both types of animals, suggesting that these changes precede symptomatic CWD cases. This observation is of potential importance in disease surveillance and diagnostics in asymptomatic animals and may eventually help prevent early spread of disease to new locations and geographical areas.

Several of the microbial taxa and metabolites found altered between CWD-positive and CWD-negative samples in our study have been previously linked to aging and disease in humans and mice, and could be physiologically relevant to CWD. This is especially interesting and promising when looking correlated changes in microbiome and metabolite levels in CWD compared to those human diseases. For instance, butyrate levels in human patients with diabetic nephropathy, which results in skeletal muscle atrophy, are significantly decreased (*36*) and a butyrate-containing diet has been shown to improve metabolism and reduce muscle atrophy in aging (*37*) and diabetic mice (*36*). This dovetails with the deceased butyrate levels in the CWD feces. GABA is a major inhibitory neurotransmitter in the brain. GABA transmission (*38*) and GABA and ornithine levels (*39*) in the cortex decrease in certain prion diseases. Our results show that the corresponding amino acid levels also decreased in the CWD feces. The secondary bile acids, deoxycholic acid (DCA) and lithocholic acid (LCA) that originate from the gut microbiome and their corresponding primary bile acids influence the neuropathology of Alzheimer’s disease (AD) and the secondary to primary bile acid ratios change significantly in AD brains (*40, 41*). While these changes have been previously reported in humans, it is possible that changes in these metabolites also occur in deer in conjunction with the change of microbiome in CWD. It was known that Eubacteriaceae (*46*) and Dorea (*47*) facilitate the conversion of primary to secondary bile acids in gut. In the CWD feces, they are significantly correlated with DCA, thus the decrease of DCA could be the results of their decrease. Both of them have no correlation with GABA and ornithine while the latter has correlation with butyrate levels in the CWD feces. Among the identified taxa at Genus and Species levels, another 2 are also known to be involved in bile salt hydrolases: Methanobrevibacter (*48*) and Enterococcus faecium (*49*). While the former have strong correlation with DCA and LCA as well as butyrate, GABA, and ornithine in the CWD feces, the latter shows no correlation. Similar to Methanobrevibacter, Olsenella also has strong correlations with DCA, LCA, butyrate, GABA, and orithine in the CWD feces. While further studies are needed to determine which of these changes are driven by factors other than CWD, these seem to be promising to explore as potential biomarkers that could serve as basis for CWD diagnostics.

It is unclear at present what is the interdependence between CWD and changes in the fecal microbiome. The changes we report here are clearly influenced by the geopgraphical origin of samples, and may be influenced by such factors as diet, seasonal changes, and other factors not included in this study. It is also possible that some of these changes reflect an overall effect on declining health in the prion-infected deer, e.g. changes in the body weight and muscle mass, wasting, or inflammation previously reported as symptoms of CWD and other prion diseases. Curiously, however, regardless of being asymptomatic and in some cases at the early stages of CWD deer still show the changes in the fecal microbiomes compared to control, as evidenced by comparison of LN and BRLN CWD-positive deer. With this knowledge, it appears especially promising to explore statistically significant changes in CWD-positive fecal microbiomes as a possible disease biomarker. Given CWD’s relatively long incubation period, this approach may prove useful for earlier detection of CWD and a substantial improvement in disease-managing strategies.

## Acknowledgements

We thank G. T. Greeshma for help with data analysis and presentation and Dr. Daniel Beiting and Dr. Lisa Mattei from the Penn Vet Center for Host-Microbial Interactions for data analysis and helpful discussions throughout the project. This work was supported by funding from the Pennsylvania Department of Agriculture and Pennsylvania Game Commission.

