## Supplemental Tables and Figures for "Prospective Fecal Microbiomics Biomarkers for Chronic Wasting Disease"

**Supplemental Table List:**

**Table S1a. Metadata for the deer samples.**

**Table S1b. Raw file names and reads information for 16S rRNA of the deer samples.**

**Table S2a. Abundance taxa count data.**

**Table S2b. Abundance clade percent data.**

**Table S3a. Alpha diversity based on taxa.**

**Table S3b. Alpha diversity based on clustered reads.**

**Tables S4. Beta diversity matrices: a-b for all; c-d for Midwest 5, Midwest 7 and South 1.**

**Tables S5. Identified biomarkers from LEfSe analysis.**

**Table S6. Filtered taxa and dentified biomarkers from Welch's t-test and Rank sum test.**

**Table S7a. Metabolomics reads data for amino acids, bile acids, and short chain fatty acids.**

**Table S7b. Metabolomics references for amino acids, bile acids, and short chain fatty acids.**

**Table S7c. Enriched taxa with significant correlation to metabolites.**

**Table S8. Identified biomarkers of taxa pairs from Welch's t-test.**

Fig. S1

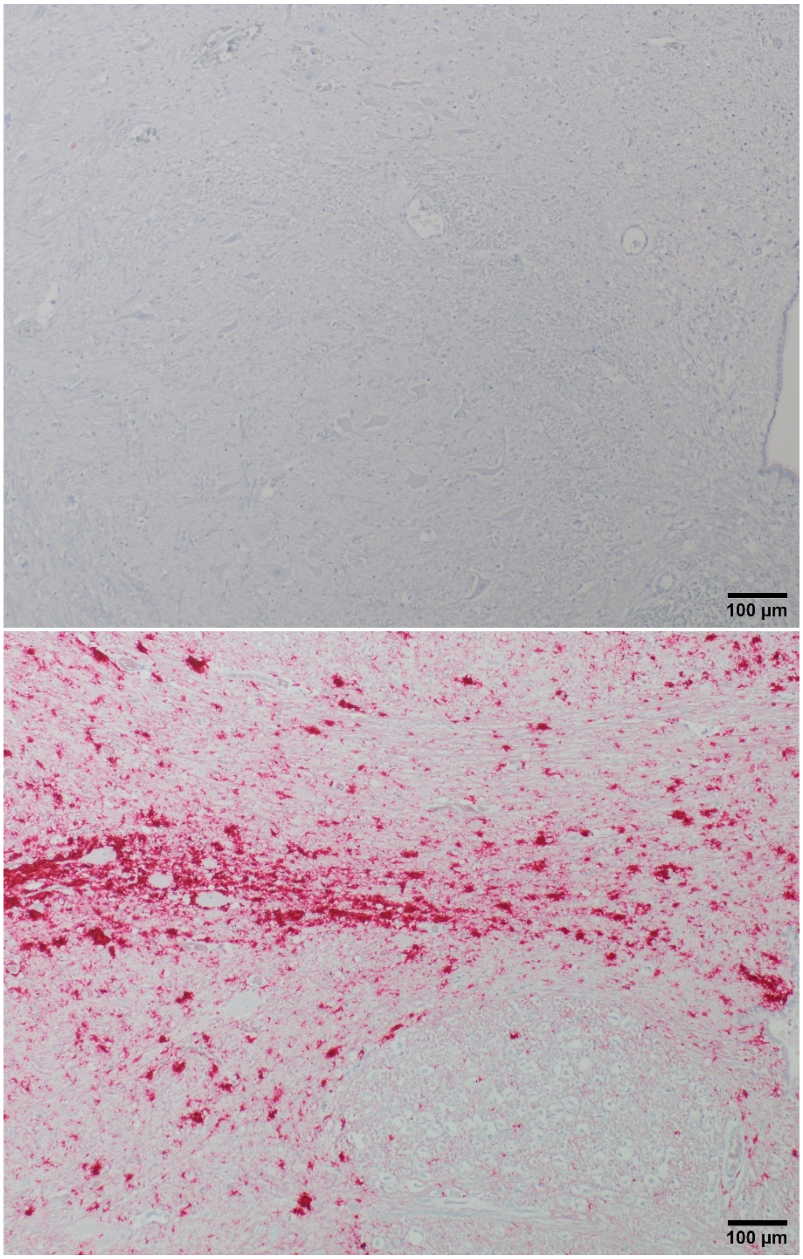


**Figure S1. Example of CWD diagnostics.** Brain section stained for prion marker are shown for a negative (top) and CWD-positive (bottom) deer sample.

Fig. S2

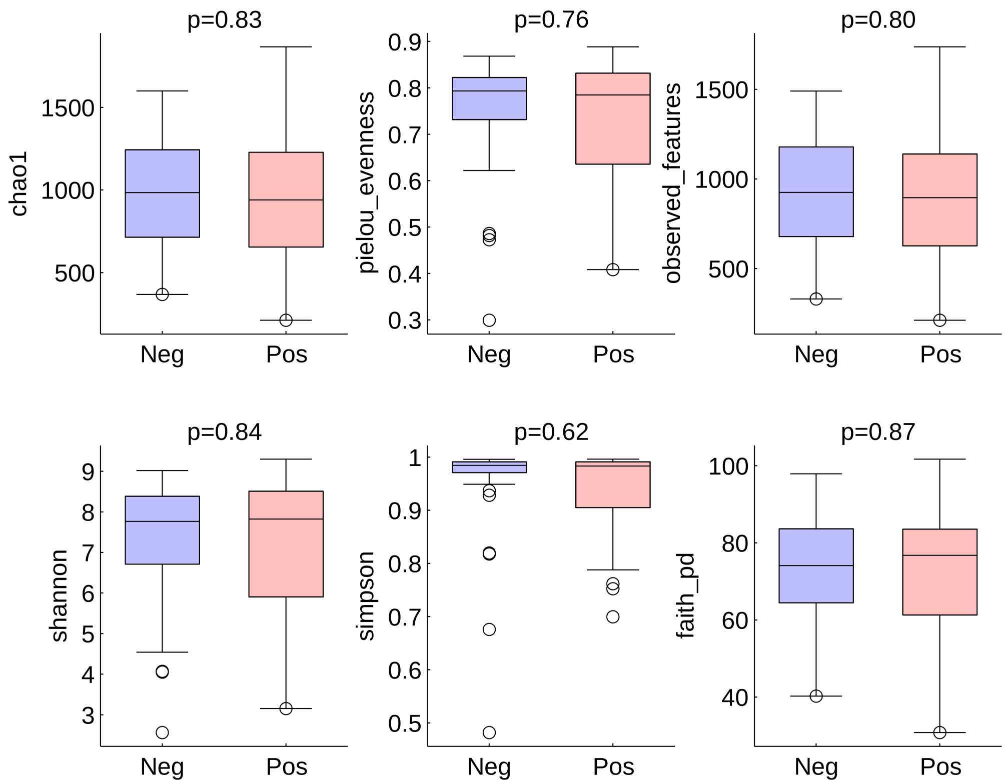


**Figure S2. Alpha diversity of the fecal samples does not depend on CWD.** Box and whisker plots showing mean and 95% confidence interval for Alpha diversity of the control (Neg) and CWD-positive (Pos) samples. P-value change is shown on the plot, n=50 in each group. Data shown in Table S3b.

Fig. S3

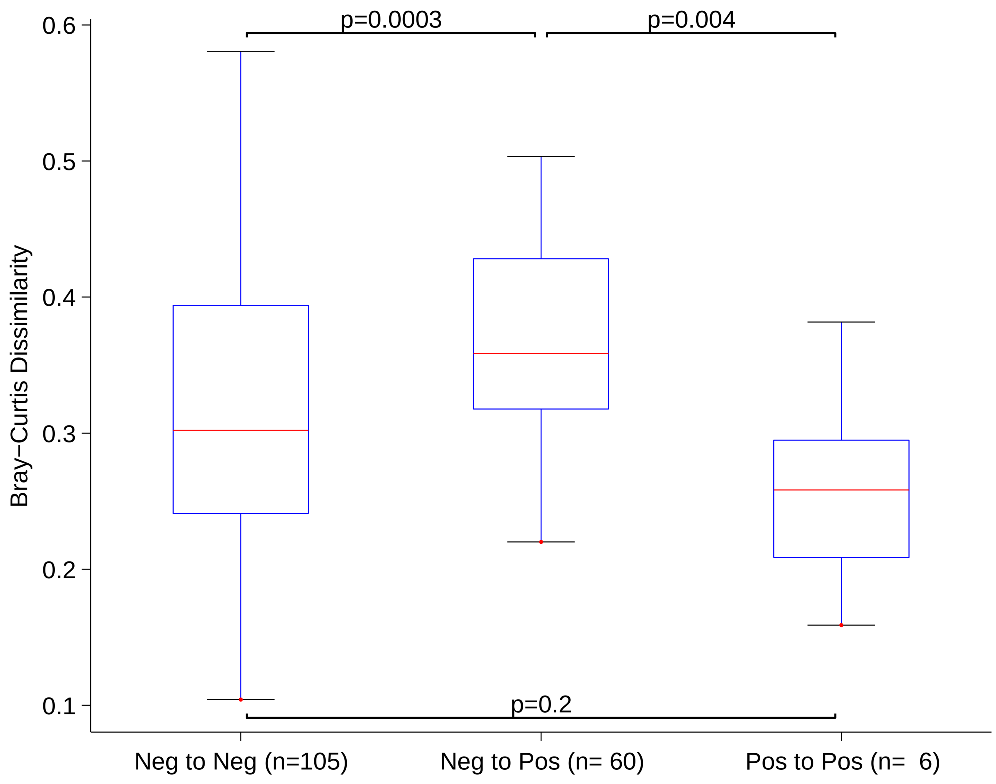


**Figure S3. Beta diversity of the fecal samples depend on CWD.** Box and whisker plots showing mean and 95% confidence interval for beta diversity of the control (Neg, n=15) and CWD-positive (Pos, n=4) samples. P-value change is shown on the plot. Data from Midwest 7 region matrix in Table S4c.

Fig. S4

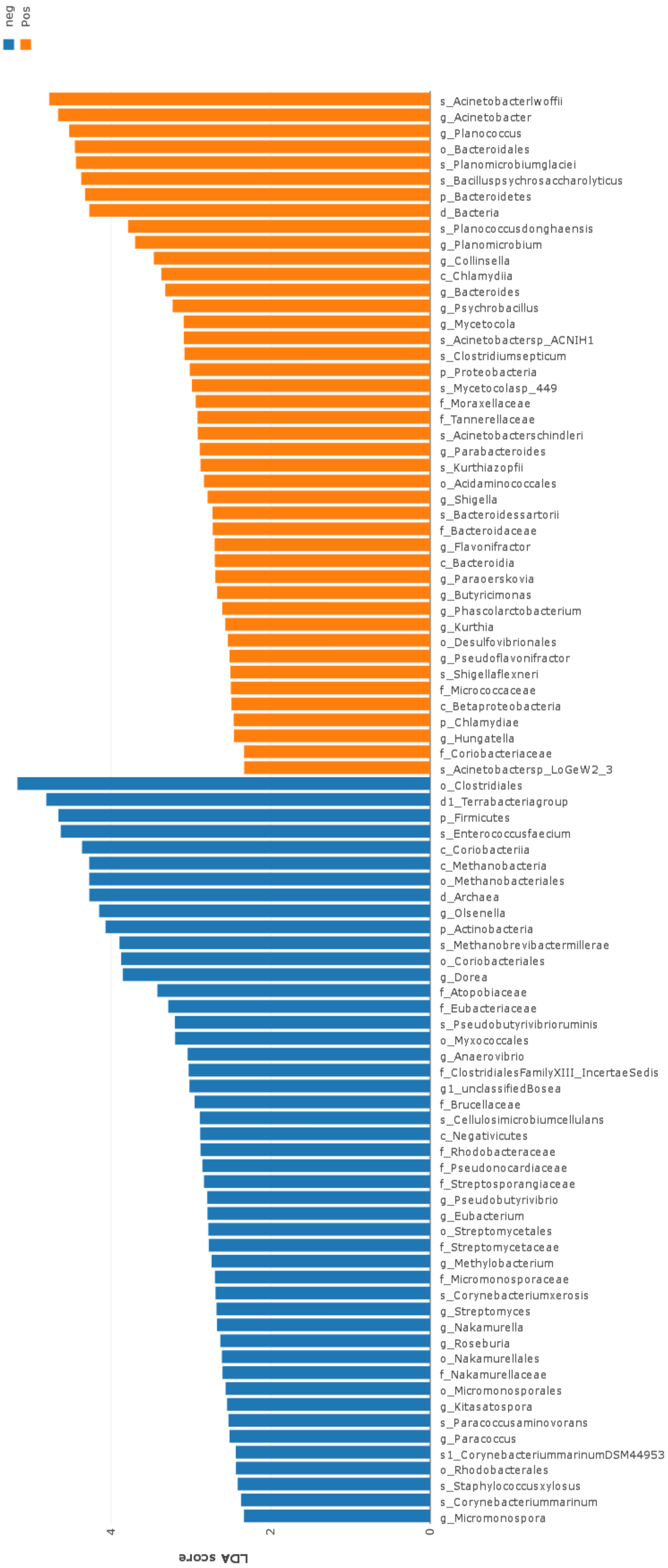


**Figure S4. LEfSe analysis result for all samples.** Bar plot shows LDA scores of taxa with p-value <0.05 for higher abundance in the CWD-negative (Neg) and CWD-positive (Pos) samples. Details shown Table S5a.

Fig. S5

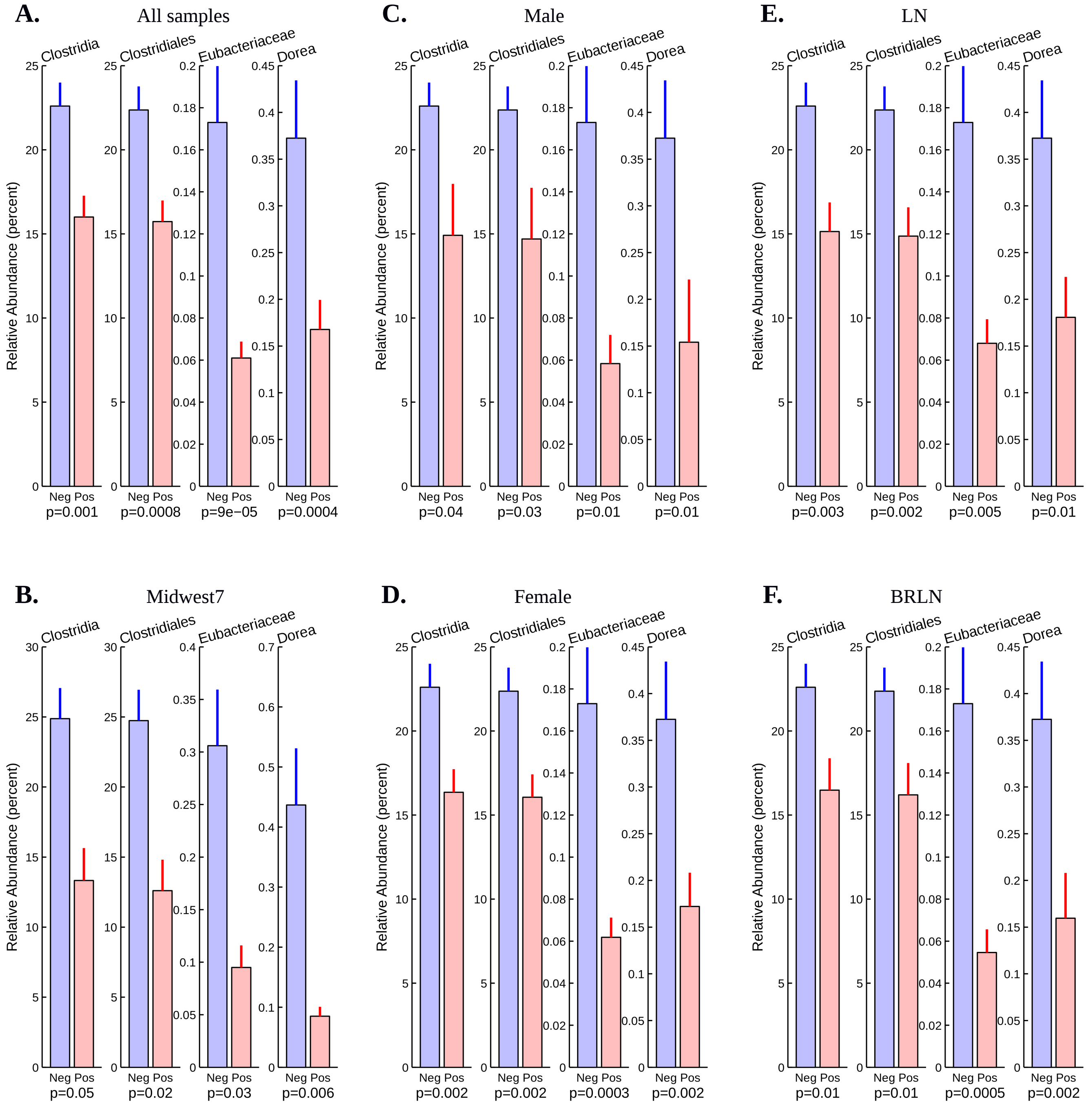


**Figure S5. Clostridia are consistently less abundant in CWD-positive samples.** CWD-positive samples show a significant reduction in abundance of the Clostridia classs, the Clostridales order, the Eubacteriacease family, and the Dorea genus. A. In all samples. B. In samples from Midwest7 region with both CWD-positive and CWD-negative deer. C. In male samples. D. In female samples. E. Positive samples with mis-folded prions in LN only. F. Positive samples with mis-folded prions in BR and LN. Bars represent geometric means. Error bars represent SEM. The p-values are from Wilcoxon rank sum test.

Fig. S6

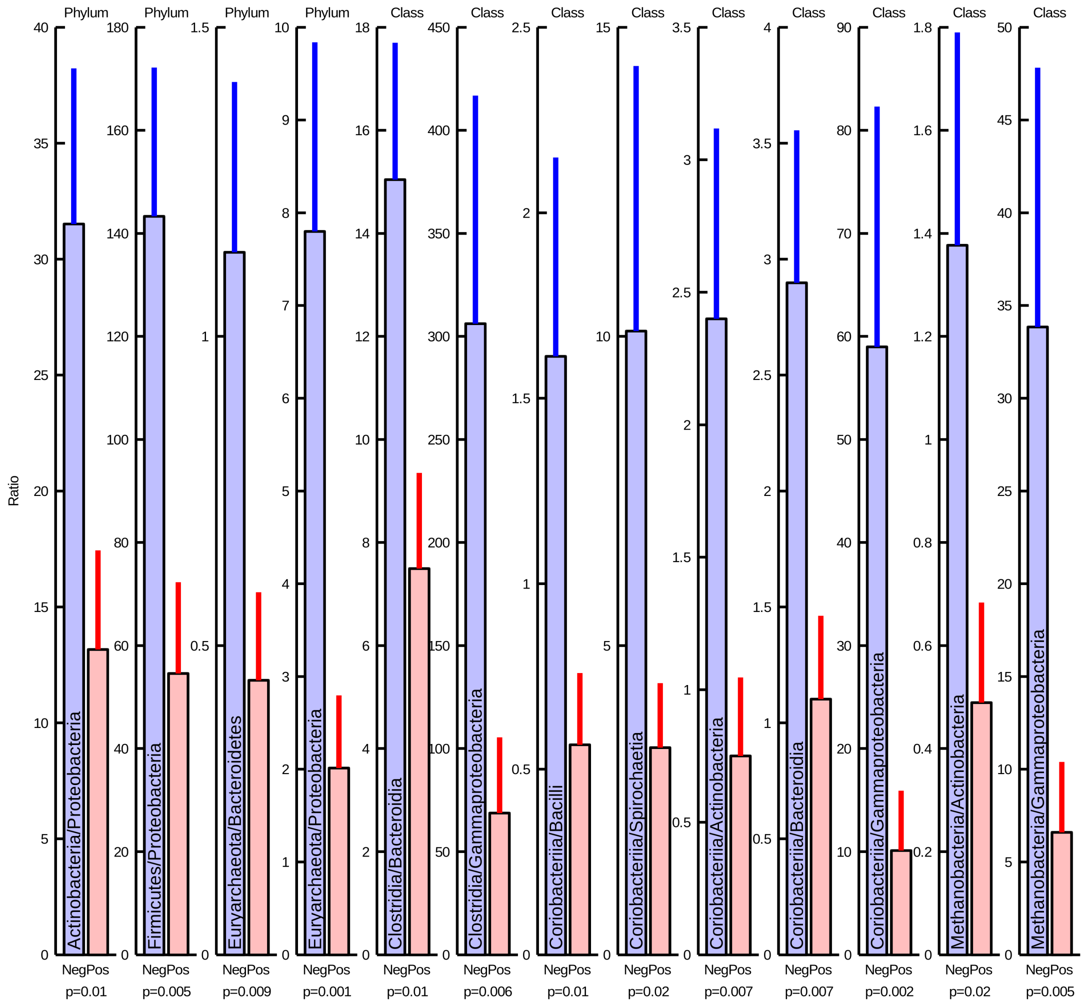


**Figure S6. Taxa ratios as potential diagnostic markers of CWD at the Phylum and Class levels.** The bar charts are for average ratios of each pairs of taxas in the control (Neg), versus CWD-positive (Pos) samples. Bars represent geometric means. Error bars represent SEM, n=50 for each set. See Table S8 for the value change in each pair.

Fig. S7

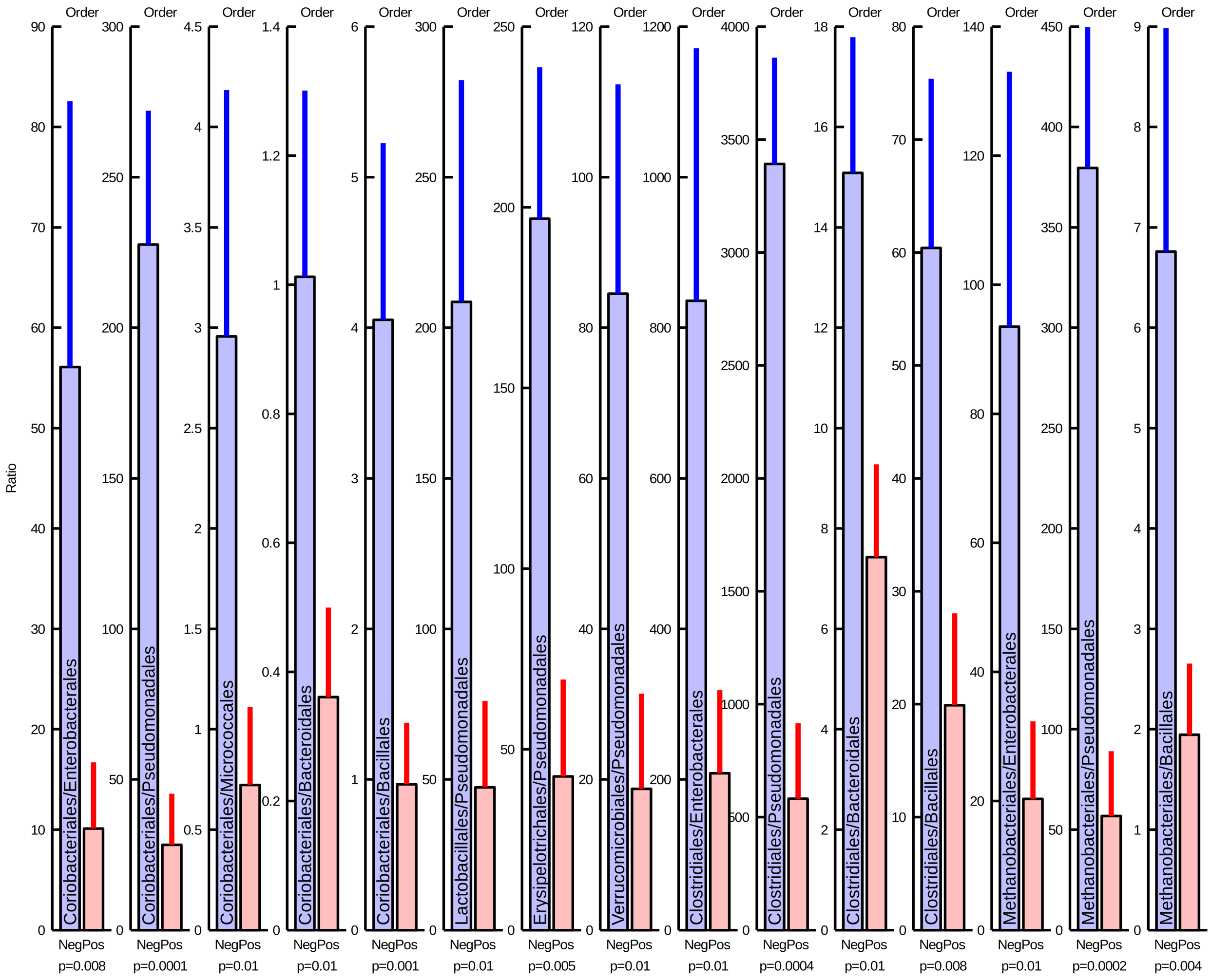


**Figure S7. Taxa ratios as potential diagnostic markers of CWD at the Order level.** The bar charts are for average ratios of each pairs of taxas in the control (Neg), versus CWD-positive (Pos) samples. Bars represent geometric means. Error bars represent SEM, n=50 for each set. See Table S8 for the value change in each pair.

Fig. S8

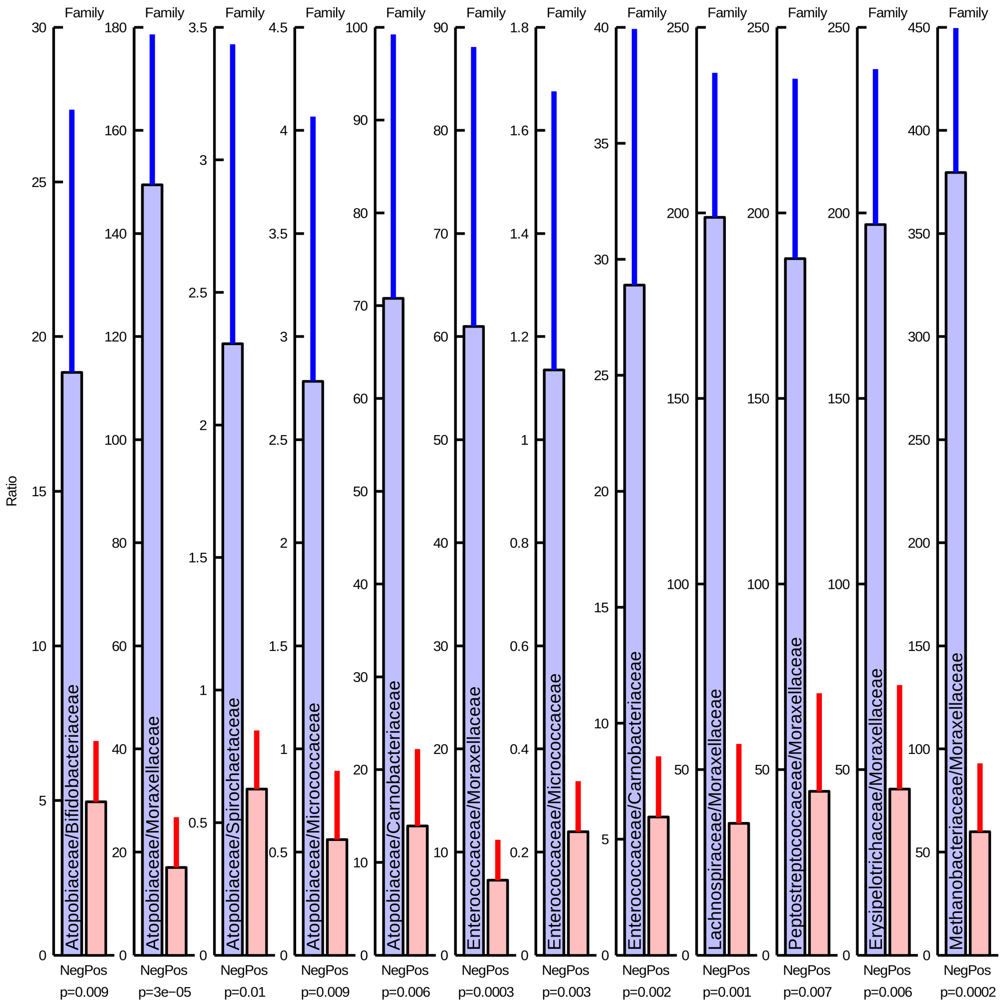


**Figure S8. Taxa ratios as potential diagnostic markers of CWD at the Family level.** The bar charts are for average ratios of each pairs of taxas in the control (Neg), versus CWD-positive (Pos) samples. Bars represent geometric means. Error bars represent SEM, n=50 for each set. See Table S8 for the value change in each pair.

Fig. S9

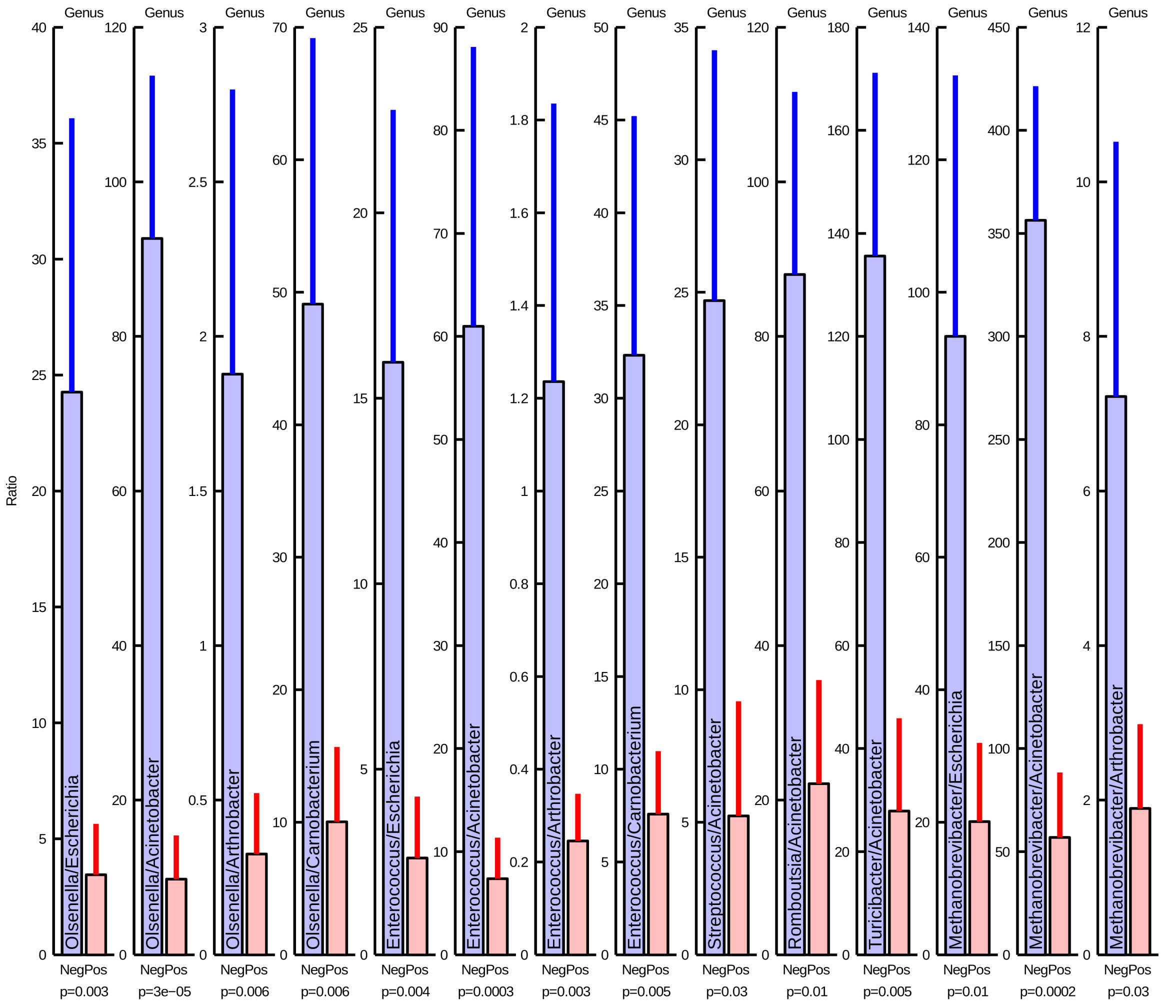


**Figure S9. Taxa ratios as potential diagnostic markers of CWD at the Genus level.** The bar charts are for average ratios of each pairs of taxas in the control (Neg), versus CWD-positive (Pos) samples. Bars represent geometric means. Error bars represent SEM, n=50 for each set. See Table S8 for the value change in each pair.

Fig. S10

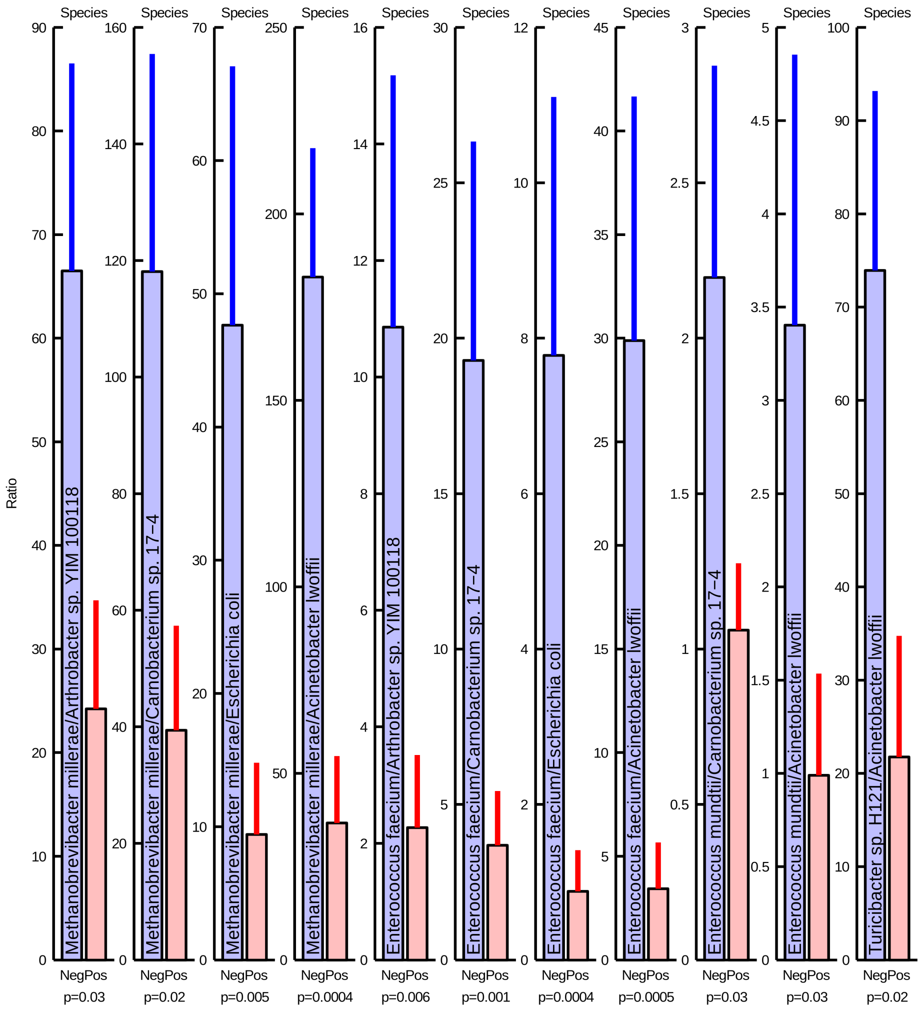


**Figure S10. Taxa ratios as potential diagnostic markers of CWD at the Species level.** The bar charts are for average ratios of each pairs of taxas in the control (Neg), versus CWD-positive (Pos) samples. Bars represent geometric means. Error bars represent SEM, n=50 for each set. See Table S8 for the value change in each pair.
